# 3D atlas of the pituitary gland of the model fish medaka

**DOI:** 10.1101/2021.05.31.446412

**Authors:** Muhammad Rahmad Royan, Khadeeja Siddique, Gergely Csucs, Maja A. Puchades, Rasoul Nourizadeh-Lillabadi, Jan G. Bjaalie, Christiaan V. Henkel, Finn-Arne Weltzien, Romain Fontaine

## Abstract

In vertebrates, the anterior pituitary plays a crucial role in regulating several essential physiological processes via the secretion of at least seven peptide hormones by different endocrine cell types. Comparative and comprehensive knowledge of the spatial distribution of those endocrine cell types is required to better understand their role during the animal life. Using medaka as the model and several combinations of multi-color fluorescence *in situ* hybridization, we present the first 3D atlas revealing the gland-wide distribution of seven endocrine cell populations: lactotropes, thyrotropes, Lh and Fsh gonadotropes, somatotropes, and *pomca*-expressing cells (corticotropes and melanotropes) in the anterior pituitary of a teleost fish. By combining *in situ* hybridization and immunofluorescence techniques, we deciphered the location of corticotropes and melanotropes within the *pomca*-expressing cell population. The 3D localization approach reveals sexual dimorphism of *tshba*- and *pomca*-expressing cells in the adult medaka pituitary. Finally, we show the existence of bi-hormonal cells co-expressing *lhb-fshb, fshb-tshba* and *lhb-sl* using single-cell transcriptomics analysis and *in situ* hybridization. This study offers a solid basis for future comparative studies of the teleost pituitary and its developmental plasticity.

**Highlights:**

- We offer the first 3D atlas of a teleost pituitary, which presents a valuable resource to the endocrinology and model fish community.
- The atlas reveals the 3D spatial distribution of the seven endocrine cell types and blood vessels in the juvenile/adult male and female pituitary.
- Gene expression for *tshba* and *pomca*, as well as the population size of cells expressing these genes, displays obvious sexual dimorphism in the adult medaka pituitary.
- Multi-color *in situ* hybridization and single cell RNA-seq reveal the existence of bi-hormonal cells, co-expressing *lhb-fshb, fshb-tshba, lhb-sl*, and a few multi-hormonal cells.
- An online version of the atlas is available at https://www.nmbu.no/go/mpg-atlas.

## Introduction

In vertebrates, the pituitary is considered the *chef d’orchestre* of the endocrine system, regulating several essential biological and physiological functions throughout the life cycle. Located beneath the hypothalamus, it is divided into the anterior part (adenohypophysis) and posterior part (neurohypophysis). The former comprises several endocrine cell types which produce and release specific peptide hormones (1, 2), controlling many important aspects of life, including growth, stress, metabolism, homeostasis, and reproduction (3).

During embryogenesis, different cellular developmental trajectories specify several endocrine cell types in the adenohypophysis, characterized by the hormones they produce (4). In general, the vertebrate adenohypophysis consists of lactotropes (producing prolactin; Prl), corticotropes (adrenocorticotropic hormone; Acth), thyrotropes (thyrotropin; Tsh), gonadotropes (follicle-stimulating and luteinizing hormone; Fsh and Lh), somatotropes (growth hormone; Gh), and melanotropes (melanocyte-stimulating hormone; α-Msh), which have specific roles in regulating certain physiological functions (2, 5). Teleosts, in addition, have an endocrine cell type that is unique to these animals, i.e. somatolactotropes (somatolactin; Sl) (6). In contrast to mammals and birds, Fsh and Lh are mostly secreted by distinct endocrine cell types in teleosts (7), although both transcripts or hormones have sometimes been observed in the same cells in some species (8–11). Unlike mammals, teleost endocrine cells are arranged in discrete zones. Lactotropes and corticotropes are commonly located in the *rostral pars distalis* (RPD), thyrotropes, gonadotropes, and somatotropes in the *proximal pars distalis* (PPD), and melanotropes and somatolactoropes in the *pars intermedia* (PI) (6, 12).

Over the past five decades, endocrine cell type organization in the teleost pituitary has been documented in various species. Despite having approximately similar patterns, the pituitary endocrine cell maps exhibit differences in terms of variety of cell types that are reported. For instance, the localization of endocrine cell populations in dorado fish shows only four distinct types of endocrine cells across the adenohypophysis (13). By contrast, the other studies describe five (Japanese medaka (14)), six (fourspine sculpin, (15); cardinal and bloodfin tetra (16)), seven (greater weever fish, (17); white seabream, (18); dimerus cichlid (19)), and eight (Atlantic halibut, (20); Nile tilapia, (21); saddle wrasse, (22)) cell types.

Even though these previous studies have provided interesting information on the spatial organization of endocrine cell populations, they lack information due to the techniques available and used at the time. First, the use of mid- and para-sagittal sections of the pituitary to reconstruct organizational patterns of endocrine cells leads to a lack of information on the lateral sides. Second, the single-labeling method and non-species specific antibodies that are typically used do not provide sufficient detail on arrangements among adjacent endocrine cell populations, or on the possible existence of multi-hormonal cells as described in mammals (23–26). These features will be important to better understand the underlying processes in fish physiology and endocrinology. Moreover, the distribution of the blood vessels within the pituitary, which play an essential role by transporting the released hormones, is poorly known. A better knowledge will help understand how endocrine cells arranged within a vascularized system that is thought to facilitate paracrine signaling in the pituitary (27). Also, since it has been shown that the pituitary is a plastic organ with changes occurring at cellular and population levels (28), it is essential to describe the cell composition, spatial organization, and vascularization of the pituitary in detail.

The Japanese medaka (*Oryzias latipes*) is a teleost model commonly used to investigate vertebrate and teleost physiology, genetics, and development, due to easy access to a wide range of genetic and molecular techniques (29, 30). We have recently used single-cell RNA sequencing to describe seven distinct endocrine cell types (expressing *prl, pomca, fshb, lhb, tshba, gh*, and *sl*) in the medaka pituitary (31). Here, we extend this study by describing differences present in the spatial distribution of the seven endocrine cell populations, in juvenile and adult fish from both sexes. Using multi-color *in situ* hybridization techniques together with single-cell transcriptomics analysis, this study offers the first 3D atlas of teleost pituitary endocrine cell populations, allowing the identification of differences in spatial distribution patterns between sexes and stages, as well as the existence of multi-hormonal cells.

## Materials and Methods

### Experimental animals

Juvenile (2-month old) and adult (6-month old) wild type medaka (WT, d-rR strain) were reared at 28 °C in a re-circulating water system (pH 7.5; 800 μS) with 14 hours light and 10 hours dark conditions. Fish were fed three times daily using artemia and artificial feed. Sex determination was based on secondary sexual characteristics (32). Experiments were conducted in accordance with recommendations on experimental animal welfare at the Norwegian University of Life Sciences.

### Quantitative Polymerase Chain Reaction (qPCR)

RNA extraction from pituitaries (n = 7) was performed as previously described in (33). Fish were euthanized by immersion in ice water and pituitaries were collected and stored at −80 °C in 300 μl of TRIzol® (Invitrogen, Carlsbad, USA) with 6 zirconium oxide beads (Bertin Technologies, Versailles, France; diameter 1.4 μm). Later, tissues were homogenized and mixed with 120 μl chloroform. The pellet was reconstituted with 14 μl of nuclease free water. Due to the size of the tissue, 3 juvenile pituitaries were pooled to represent one replicate of each sample. A total of 33 ng of RNA was used to synthesize cDNA using SuperScript III Reverse Transcriptase (Invitrogen, Carlsbad, CA, USA) and random hexamer primers (Thermofisher scientific). 5× diluted cDNA samples were analyzed in duplicate, using 3 μL of the cDNA and 5 μM each of forward and reverse primer in a total volume of 10 μL (**Table 1**). The parameter cycle was 10 min pre-incubation at 95 °C, followed by 42 cycles of 95 °C for 10 s, 60 °C for 10 s and 72 °C for 6 s, followed by melting curve analysis to assess PCR product specificity. The mRNA level was normalized using *rpl7* as reference gene as it was found no significant difference of expression between groups.

**Table 1.**
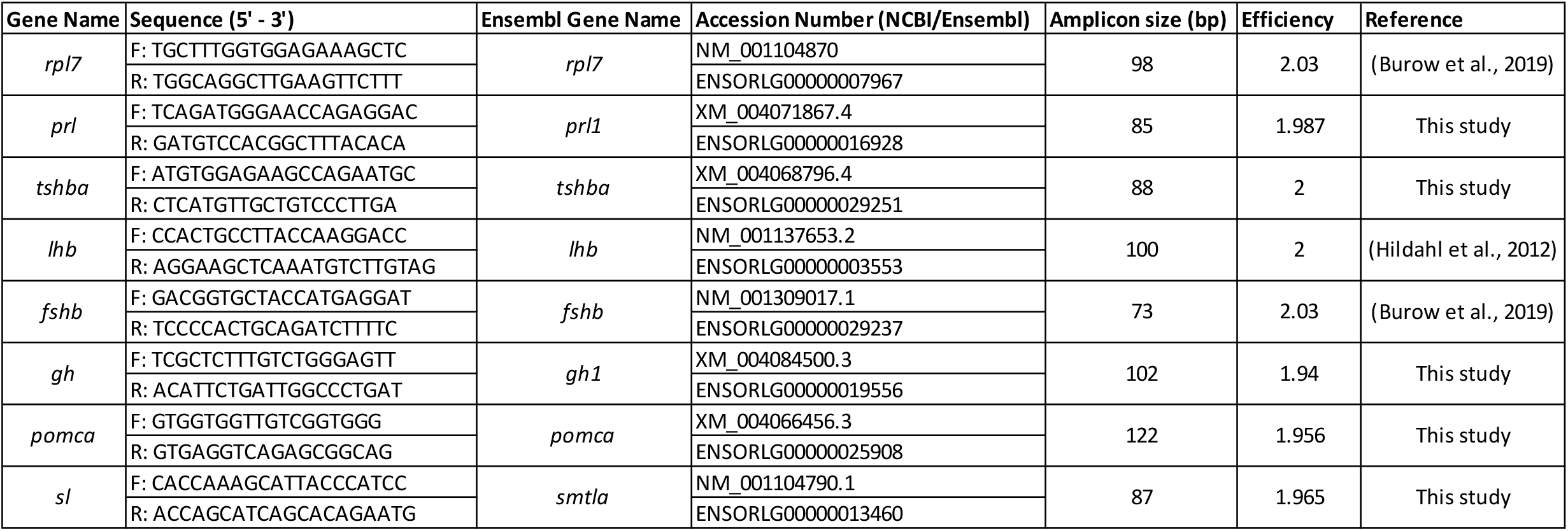
Primer sequences used for the mRNA level analysis in the medaka pituitary.

### Multi-color fluorescence *in situ* hybridization (Multi-color FISH)

#### Tissue preparation

Fish were euthanized by immersion in ice water. Brain and pituitary were taken and fixated overnight at 4 °C in 4% paraformaldehyde (PFA, Electron Microscopy Sciences, Hatfield, Pennsylvania) diluted with phosphate buffered saline with Tween (PBST: PBS, 0.1%; Tween-20), approximately 50× the tissue volume. Tissue was then dehydrated using a series of increasing ethanol concentrations: 25%, 50%, 75%, 96% followed by a storage in 100% methanol at −20 °C until use.

#### Cloning and RNA probe synthesis

DNA sequences for the probes were obtained from NCBI as listed in **Table 2**. Sequences are selected according to the high expression in the pituitary for those having more than one paralog in the medaka genome (*tshb* and *pomc*). PCR primers for the amplification of the probe genes were designed from transcribed sequences (mRNA) for each gene using Primer3 (https://primer3.ut.ee/). Following RNA extraction and cDNA synthesis as described above, cDNA was used to amplify the sequence of interest by PCR using Taq DNA polymerase (Thermo Fisher Scientific) with a 3-min denaturation step at 94 °C, followed by 35 cycles at 94 °C for 15s, 50 °C for 15s, and 72 °C for 60s, and finally 1 cycle of 72 °C for 5 mins. The amplified PCR products were isolated using a gel extraction kit (Qiagen) and cloned into the pGEM-T Easy vector (Promega) following manufacturer instructions and verified by sequencing. PCR products from the verified plasmids were used as template to synthetize sense and anti-sense complementary RNA probes using *in vitro* transcription with T7 or SP6 RNA polymerase (Promega, Madison, Wisconsin). RNA probes were tagged with dinitrophenol-11-UTP (DNP, Perkin Elmer, Waltham, Massachusetts), fluorescein-12-UTP (FITC, Roche Diagnostics), or digoxigenin-11-UTP (DIG, Roche Diagnostics). Finally, the probes were purified using the Nucleospin RNA clean-up kit (Macherey-Nagel, Hoerdt, France) and the concentration was measured using the Epoch Spectrophotometer System (BioTek, Winooski, VT, USA).

**Table 2.**
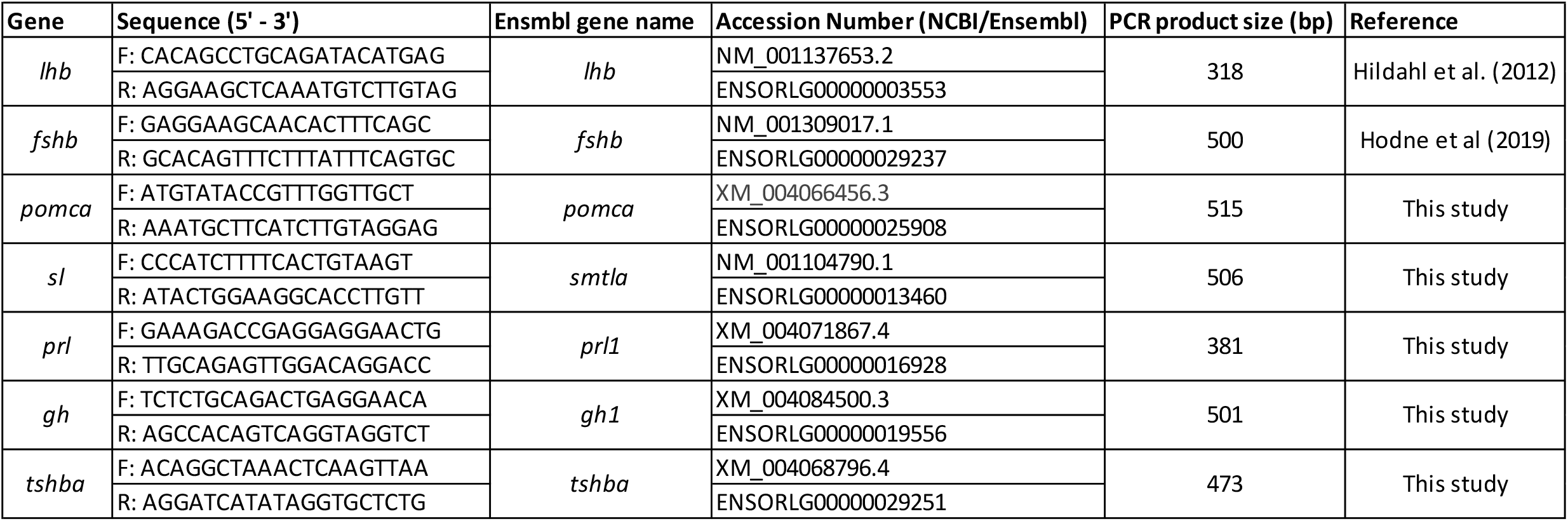
Primer sequences used to make the *in situ* hybridization (ISH) probes of seven endocrine cell types in the medaka pituitary.

#### Multi-color fluorescence *in situ* hybridization (FISH)

Multi-color FISH was performed as previously described in (34) with minor modifications. Tissues were serially rehydrated, and the pituitary was detached from the brain. Afterwards, whole pituitaries were hybridized with the probes (0.11 – 3.17 ng/μl) for 18 hours at 55 °C, and incubated with different combinations of anti-DNP- (Perkin Elmer), anti-FITC-, and anti-DIG-conjugated antibodies (Roche Diagnostics), followed by TAMRA- (Thermofisher), Cy5- (Perkin Elmer) and FITC-conjugated tyramides (Sigma). The nuclei were stained with DAPI (1:1000, 4’, 6-diamidino-2-phenylindole dihydrochloride; Sigma). The absence of labeling when using sense probes was used to confirm the specificity of the anti-sense probes. Whole pituitaries were mounted using Vectashield H-1000 Mounting Medium (Vector, Eurobio/Abcys) between microscope slides and cover slips (Menzel Glässer, VWR) with spacers (Reinforcement rings, Herma) in between for the juveniles, and between two cover slips with spacers for adults.

### Combined FISH and Immunofluorescence (IF)

To distinguish the localization of adrenocorticotropic releasing hormone (Acth) and alpha-melanocyte stimulating hormone (α-Msh) cells within *pomca*-expressing cells separately, IF was performed using the antibodies shown in **Table 3**. After FISH for *pomca* labelled with FITC-conjugated tyramide, the pituitaries were embedded in 3% agarose (H_2_O) and para-sagittally sectioned with 60 μm thickness using a vibratome (Leica). From a single pituitary, odd and even ordered slices were processed to detect Acth and α-Msh IF, respectively. Tissue slices were incubated for 10 minutes at room temperature (RT) in permeabilizing buffer (0.3 % Triton in PBST) with agitation, before incubation for 1 hour at RT in blocking solution (Acth: 3% normal goat serum (NGS); 0.3% Triton; 1% dimethylsulfoxide (DMSO) in PBST; α-Msh: 3% NGS; 5% Triton; 7% DMSO in PBST). Sections were then incubated at 4 °C overnight with primary antibodies (diluted in blocking) or without (control), followed by 4 hours at RT with secondary antibodies (diluted in blocking) with extensive PBST washes in between. Nuclei were stained with DAPI (1/1000). Controls were performed as described above incubating the tissue without the primary antibodies and confirming the absence of fluorescent signals. The antibody dilution is provided in **Table 3**.

**Table 3.**
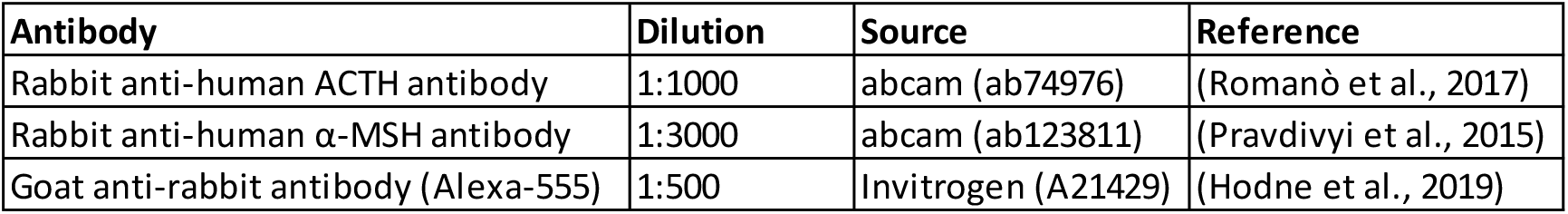
Primary and secondary antibodies used for immunofluorescence (IF) to distinguish Acth and α-Msh cells from *pomca*-expressing cells in the medaka pituitary.

| Antibody | Dilution | Source | Reference |
| --- | --- | --- | --- |
| Rabbit anti-human ACTH antibody | 1:1000 | abcam (ab74976) | (Romanò et al., 2017) |
| Rabbit anti-human α-MSH antibody | 1:3000 | abcam (ab123811) | (Pravdivyi et al., 2015) |
| Goat anti-rabbit antibody (Alexa-555) | 1:500 | Invitrogen (A21429) | (Hodne et al., 2019) |

### Blood vessel staining

Blood vessels were stained by cardiac perfusion as previously described in (35). The fish were anesthetized with 0.04% Tricaine (pH 7), and the anterior abdomen was cut to let the heart opening adequately wide for the injection. Afterwards, 0.05% of DiI (1,1′-Dioctadecyl-3,3,3′,3′-Tetramethylindocarbocyanine Perchlorate; Invitrogen) solution diluted in 4% PFA (in PBS) was administered to the *bulbus arteriosus* through the ventricle using a glass needle. The pituitary was dissected and fixated in 4% PFA (in PBS) for 2 hours in the dark before being washed 2 times with PBS and mounted as described above.

### Image processing and analysis

Fluorescent images were obtained using an LSM710 Confocal Microscope (Zeiss, Leica) with 25× (for adult pituitary) and 40× (for juvenile pituitary) objectives. Filters with wavelength of 405 (DAPI), 555 (TAMRA; Alexa-555), 633 (Cy5) and 488 (FITC; Alexa-488) nm were used. Due to the size of the adult pituitaries, the image acquisition was done from the dorsal and ventral sides of the pituitary with some overlaps in the middle. In conjunction with the microscope, ZEN software (v2009, Zeiss) was used to process the images, and ImageJ (1.52p; http://rsbweb.nih.gov/ij/) was used for processing z-projections from confocal image stacks. The dorsal and ventral stacks of adult pituitaries were aligned using HWada (https://signaling.riken.jp/en/en-tools/imagej/635/) and StackReg plugin (http://bigwww.epfl.ch/thevenaz/stackreg/), before presenting them in orthogonal views.

### 3D atlasing

While juvenile pituitaries were imaged as one block, adult pituitaries were imaged from the ventral and dorsal side with confocal imaging as described above. The two sides of the adult pituitaries were then merged using landmarks as visible with the DAPI staining. Finally, eight pituitaries labeled for different markers were aligned to the same coordinate system also using manually selected landmarks. These data were used for the creation of four 3D atlases of the pituitary gland, using the principle approaches outlined in (36).

The creation of the 3D atlases involved several steps. Merging and alignment was done using LandmarkReg (https://github.com/Tevemadar/LandmarkReg, with accompanying utilities https://github.com/Tevemadar/LandmarkReg-utils). Image stacks were saved in NIfTI format (https://imagej.nih.gov/ij/plugins/nifti.html) and converted with the “NIfTI2TopCubes” utility before the matching anatomical positions (“landmarks”) were manually identified in volume-pairs. Both the signal from endocrine cells and the DAPI background were inspected, and four or more landmarks were recorded for each volume-pair. In case of ventral-dorsal half images (adult samples), a custom utility “PituBuild” was used for merging the two halves, based on partial overlap. Finally, each set of complete pituitary volumes was aligned to a common anatomical space using the “Match” utility. The resulting NIfTI volumes were then converted to TIFF stacks for viewing and analysis.

To enable 3D viewing of the pituitary atlases, data were prepared for the MeshView tool (RRID:SCR_017222, https://www.nitrc.org/projects/meshview/) developed for 3D brain atlas viewing (37). Volumes were first binarized using the “BinX” utility with a threshold value of 50. MeshGen (https://www.nitrc.org/projects/meshgen/) was then used to generate surface meshes in standard STL format (http://paulbourke.net/dataformats/stl/) before convertion using PackSTL (https://github.com/Tevemadar/MeshView-PackSTL) to allow viewing in MeshView.

### Single cell transcriptomics analysis (scRNA-seq)

We used processed scRNA-seq dataset for male and female pituitaries (31) and filtered out red blood cells to avoid noise. Next, we applied a cut-off to differentiate between cells with high and low expression for specific genes (Supp. Fig. 1). This dataset is further used to generate the pair-wise scatterplots using the R package ggplot2 (version 2_3.3.2), to show cells expressing more than one hormone-producing gene. Finally, we got 191 and 229 multiple hormone-producing cells in female and male pituitaries, respectively. We generated clustered heatmaps using pheatmap (version 1.0.12) to visualize the expression levels of hormone-producing genes in each cell expressing multiple hormones.

### Statistical analysis

Levene’s test was performed to analyze the homogeneity of variance while the normality was tested using the Saphiro-Wilk Normality test. The differences in mRNA levels were evaluated using One-way ANOVA followed by Tukey *post hoc* test. The data are shown as mean + SEM (Standard Error of Mean) unless otherwise stated in the figure legend. *p* <0.05 was used as a threshold for statistical significance.

### Data availability

All image files are available in a data repository (https://doi.org/10.18710/NOGJQ2). The 3D atlases (Fig. 1) can be found on a webpage containing explanatory videos and other types of data completing the online pituitary atlas (https://www.nmbu.no/go/mpg-atlas), allowing easy access and navigation through the data.

**Figure 1.**
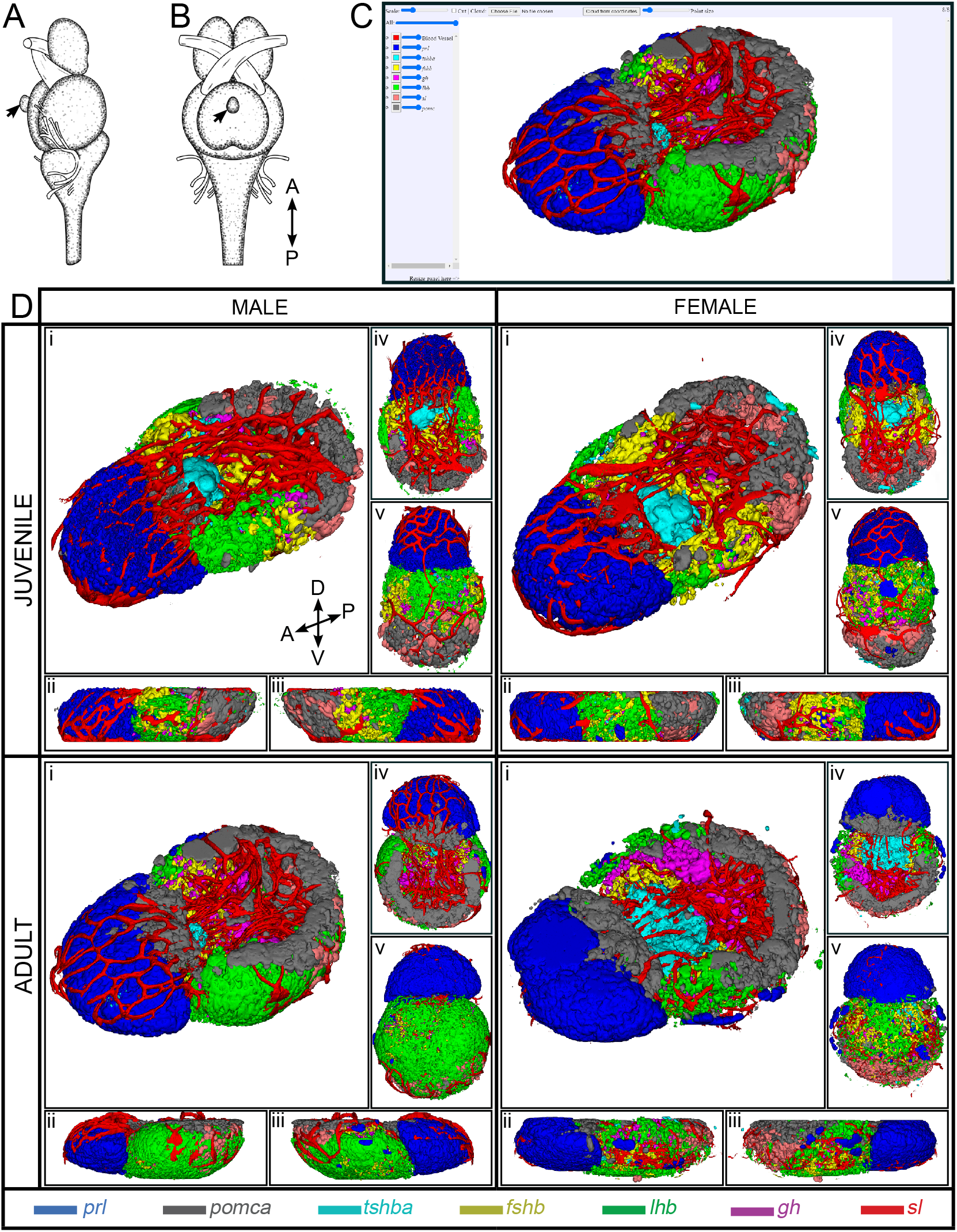
3D reconstruction of medaka anterior pituitary containing seven endocrine cell populations. Illustration of brain and pituitary (pointed by black arrow) of medaka from lateral view (A) and ventral view (B) (Figure is adopted and modified with permission from (63)). The navigation platform of 3D spatial distribution endocrine cell population that are now available online (C). Snapshots of 3D reconstruction of endocrine cell population from juvenile and adult medaka male and female (D). The snapshots were captured from different perspectives: free viewpoint (i), lateral (ii and iii), dorsal (iv) and ventral (iv). Four direction arrows display the direction of the pituitary (A: anterior; P: posterior; D: dorsal; V: ventral). The color legend shows the color code for each cell type.

## Results

### 3D atlases of the medaka pituitary

Using several combinations of multi-color FISH, we combined the labeling to form four 3D pituitary atlases now available online (http://meshview.uiocloud.no/Medaka-pituitary/?JF; http://meshview.uiocloud.no/Medaka-pituitary/?JM; http://meshview.uiocloud.no/Medaka-pituitary/?AF; http://meshview.uiocloud.no/Medaka-pituitary/?AM), allowing us to precisely localize the seven endocrine cell types (*prl, pomca, tshba, fshb, lhb, gh* and *sl*) in the adenohypophysis of medaka.

#### 1. *prl*-expressing cells (lactotropes)

In both adults and juveniles (Supp. Fig. 2), *prl*-expressing cells make up almost the entirety of the RPD from the dorsal to the ventral side of the pituitary, without any obvious difference in localization pattern between males and females. They border on and intermingle with a *pomca*-expressing cell population (Supp. Fig. 9). In some fish, a few *prl*-expressing cells are also localized peripherally in the dorsal area of PPD (data not shown).

#### 2. *pomca*-expressing cells (corticotropes and melanotropes)

*pomca*-expressing cells are observed in two distinct places in the pituitary. One population is localized in the dorsal part of RPD, where it is mostly clustered in the middle if observed from transverse point of view, and the second is detected in the PI area (Supp. Fig. 3). While the first one is adjacent to and mixing with *prl*-expressing cells, in close proximity to *tshba*-expressing cells (Supp. Fig. 9), the second population intermingles with *sl*-expressing cells (Supp. Fig. 10).

#### 3. *tshba*-expressing cells (thyrotropes)

*tshba*-expressing cells are localized in the dorsal side of anterior PPD towards the PN, next to the *prl*- and *pomca*-expressing cells (Supp. Fig. 4 and Supp. Fig. 9). From a transverse point of view, *tshb*-expressing cells are mostly concentrated in the middle part of the PPD (Supp. Fig. 4) where they border and mix with *fshb*-expressing cells (Supp. Fig. 11).

#### 4. *fshb*-expressing cells (gonadotropes)

*fshb*-expressing cells are detected from the anterior to middle part of the PPD, distributed in both lateral sides of the pituitary from a transverse point of view (Supp. Fig. 5). These cells cover the PN from a frontal point of view (Supp. Fig. 5). They border and mix with *tshba*-expressing cells in the dorsal (Supp. Fig. 11), *lhb*-expressing cells in the ventral (Supp. Fig. 12) and *gh*-expressing cells in the posterior part of the PPD (Supp. Fig. 13).

#### 5. *lhb*-expressing cells (gonadotropes)

In both juveniles and adults, *lhb*-expressing cells are commonly distributed in the peripheral area of the PPD, covering almost the entire ventral side of the pituitary (Supp. Fig. 6). In adults, *lhb*-expressing cells are also localized in the proximity of peripheral area of the PI (Supp. Fig. 6). These cells border and mix with *fshb*-expressing cells in the PPD (Supp. Fig. 12), and with *pomca*- and *sl*-expressing cells in the PI of adult pituitary (Supp. Fig. 10).

#### 6. *gh*-expressing cells (somatotropes)

*gh*-expressing cells are localized on the dorsal side of the PPD towards the PN area (Supp. Fig. 7). Similar to *fshb*-expressing cells, which are distributed to both lateral sides of the pituitary, *gh*-expressing cells are more towards the posterior part of the PPD, encompassing and bordering with the PN (Supp. Fig. 7), mixing with *fshb*-expressing cells (Supp. Fig. 13).

#### 7. *sl*-expressing cells (somatolactotropes)

*sl*-expressing cells are intermingled within *pomca*-expressing cells located in the PI (Supp. Fig. 8 and Supp. Fig. 10). In the adult pituitary, these cells border and mix with *lhb*-expressing cells that are detected in the proximity of the PI (Supp. Fig. 10).

#### 8. Distinction of Acth and α-Msh cell populations

The combination of FISH for *pomca* with IF for Acth or α-Msh shows that that Acth cells overlap the entire *pomca* signal, while melanotropes overlap *pomca* signals that are located in the PI, both in adults (Fig. 2) and in juvenile pituitaries (Supp. Fig. 14).

**Figure 2.**
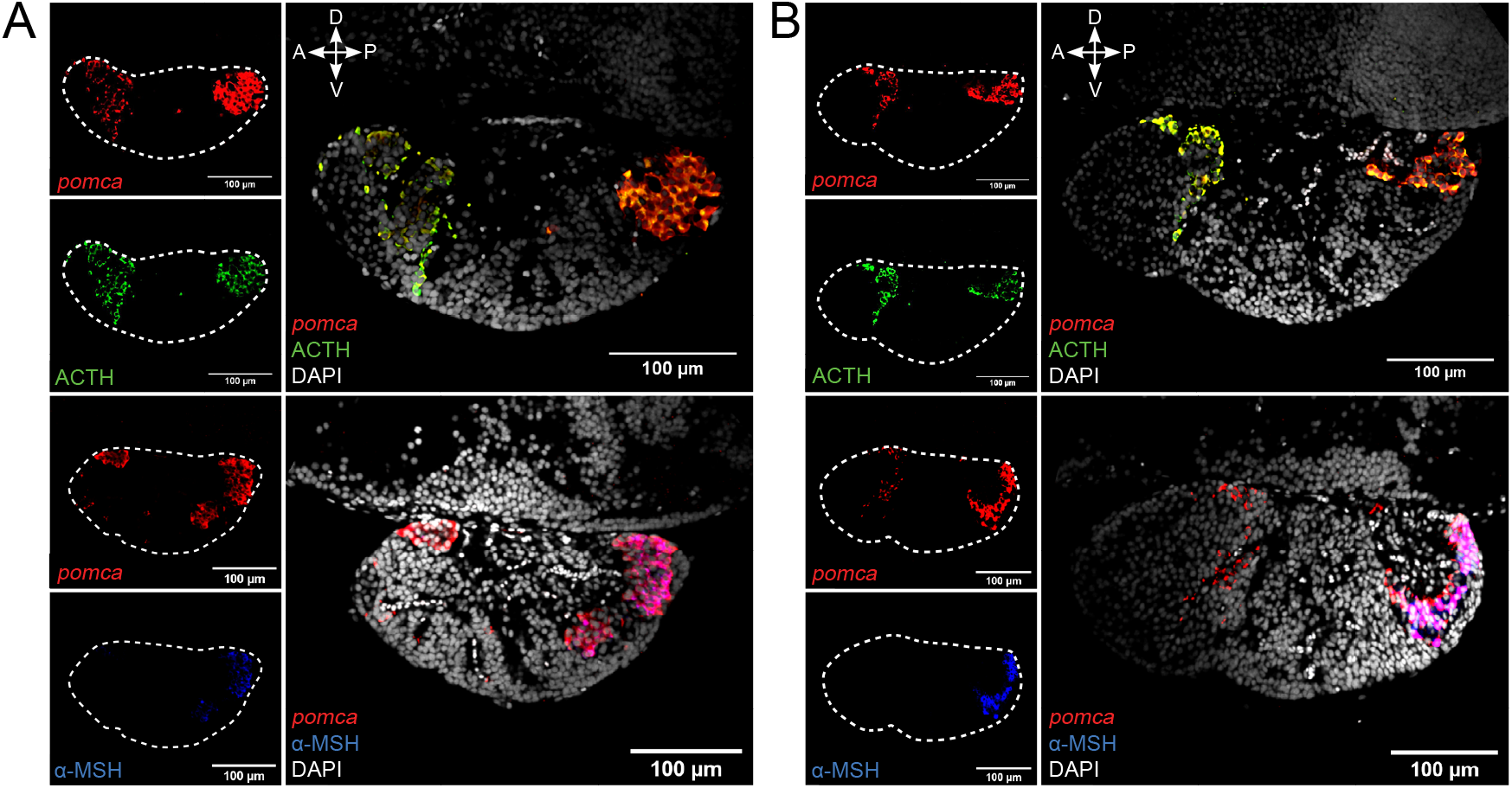
The combination of FISH for *pomca* and IF for Acth or α-Msh allows the distinction of two clear *pomca* expressing cell populations. The distinction of Acth (green) and α-Msh (blue) producing cells from *pomca*-labelled (red) in the pituitary from adult male and female medaka. Dashed line represents the pituitary as shown in the right panel. Four direction arrows display the direction of the pituitary (A: anterior; P: posterior; D: dorsal; V: ventral).

#### 9. Blood vessels

3D reconstruction shows that blood vessels encompass the entire adenohypophysis, without any obvious difference observed between sexes and stages (Fig. 3).

**Figure 3.**
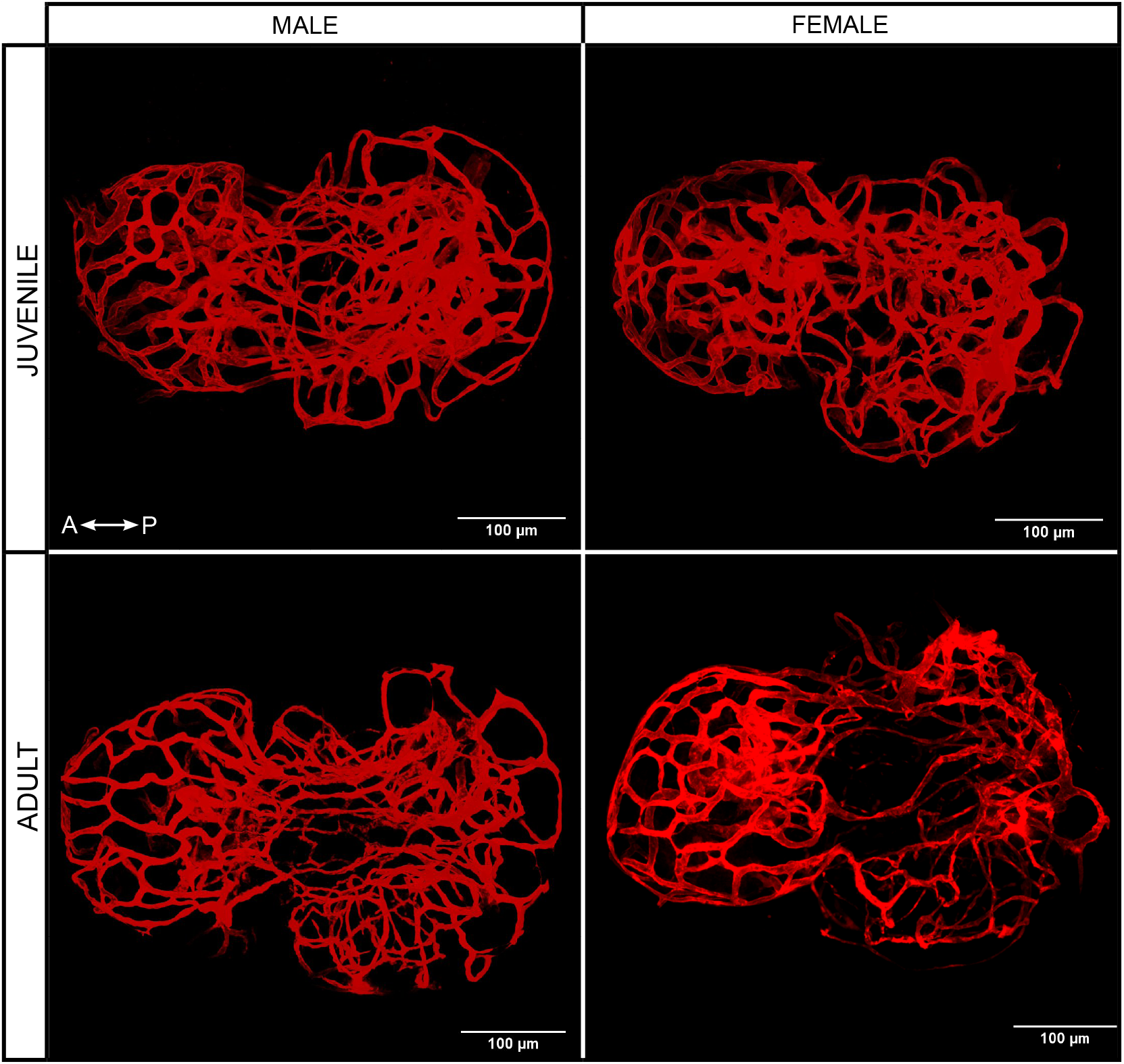
3D projection of blood vessels from juvenile and adult male and female medaka pituitary from the dorsal side. Left right arrow symbol shows the direction of the pituitary (A: anterior; P: posterior).

### Sex and stage differences

FISH and qPCR data did not reveal any difference in *prl*- and *gh*-expressing cell number or mRNA levels between sexes and stages. In contrast, adult females show significantly higher mRNA levels in both *fshb* (*p* < 0.01) and *lhb* (*p* < 0.0001) compared to the other groups, while *lhb* mRNA levels are significantly higher in adult males than in juveniles (*p* < 0.05) (Fig. 4), despite no obvious difference in the cell population size (Supp. Fig. 4-5). *sl* mRNA levels are significantly higher in juveniles compared to adult males (*p* < 0.05) (Fig. 4) without any obvious difference in cell number (Supp. Fig. 8).

**Figure 4.**
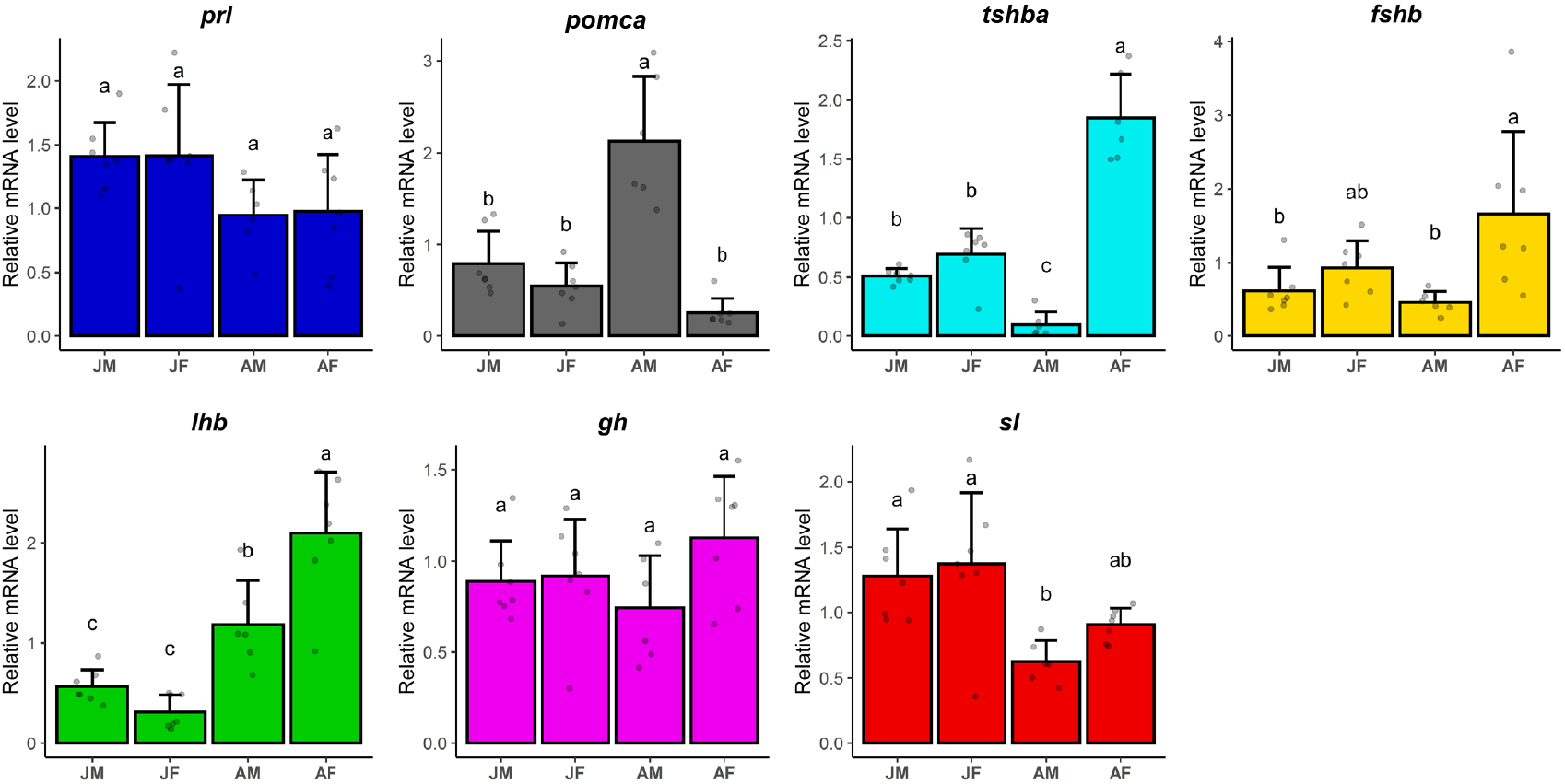
Relative mRNA levels of endocrine cell markers (*prl, tshba, fshb, lhb, gh, sl* and *pomca*) in juvenile and adult medaka males and females (juvenile male, JM; juvenile female, JF; adult male, AM; adult female, AF). Graphs are provided as mean + SEM, and jitter dots represent N. Different letters display statistical differences (*p* < 0.05) between groups as evaluated by One-way ANOVA followed by Tukey *post hoc* test.

In contrast, FISH revealed that the population of *tshba*-expressing cells is larger in adult females, expanding from the anterior to the middle part of PPD, whereas in adult males and juveniles they are found only in the anterior part of the PPD (Supp. Fig. 4). This agrees with the significantly higher *tshba* mRNA levels observed by qPCR in the adult female group compared to the other groups (*p* < 0.0001) (Fig. 4). The *tshba*-expressing cell population also appears larger in juveniles than in adult males using FISH, in line with significantly higher level of *tshba* mRNA observed by qPCR in juveniles compared to in adult males (*p* < 0.05) (Fig. 4 and Supp. Fig. 4).

*pomca* mRNA levels in adult male are significantly higher than the other groups (*p* < 0.0001), and this agrees with the *pomca*-expressing cell population illustrated in Fig. 2, Supp. Fig. 3 and Supp. Fig. 14, where there are bigger populations of *pomca* cells in adult male than the other groups.

Despite obvious differences in mRNA transcript levels and cell number observed with qPCR and FISH/IF labeling, we could not observe any difference in the proportion of each cell type between males and females with the scRNA-seq data.

### Cells producing multiple hormones

Using scRNA-seq data, we observed some bi-hormonal cells in the pituitary of adult medaka (Fig. 5). Both sexes show a number of cells co-expressing *lhb*- and *fshb*-, *lhb*- and *tshba*- and *fshb* and *tshba*. Meanwhile, some cells co-expressing *lhb* and *sl, fshb* and *sl, fshb* and *prl, fshb* and *pomca, tshba* and *prl, tshba* and *pomca, prl* and *gh*, and *prl* and *pomca* are unique to adult males, whereas co-expression of *fshb*- and *gh*-expressing cells is only found in adult females (Fig. 5A). The co-expression between *lhb-fshb* and *fshb-tshba* in both sexes and *lhb-sl* in adult male was confirmed using multi-color FISH (Fig. 6). While the co-localization of cells expressing *lhb-fshb* and *fshb-tshba* was observed in several individuals, the *lhb-sl* expressing cells were observed only in 1 out of 13 adult male pituitaries analyzed.

**Figure 5.**
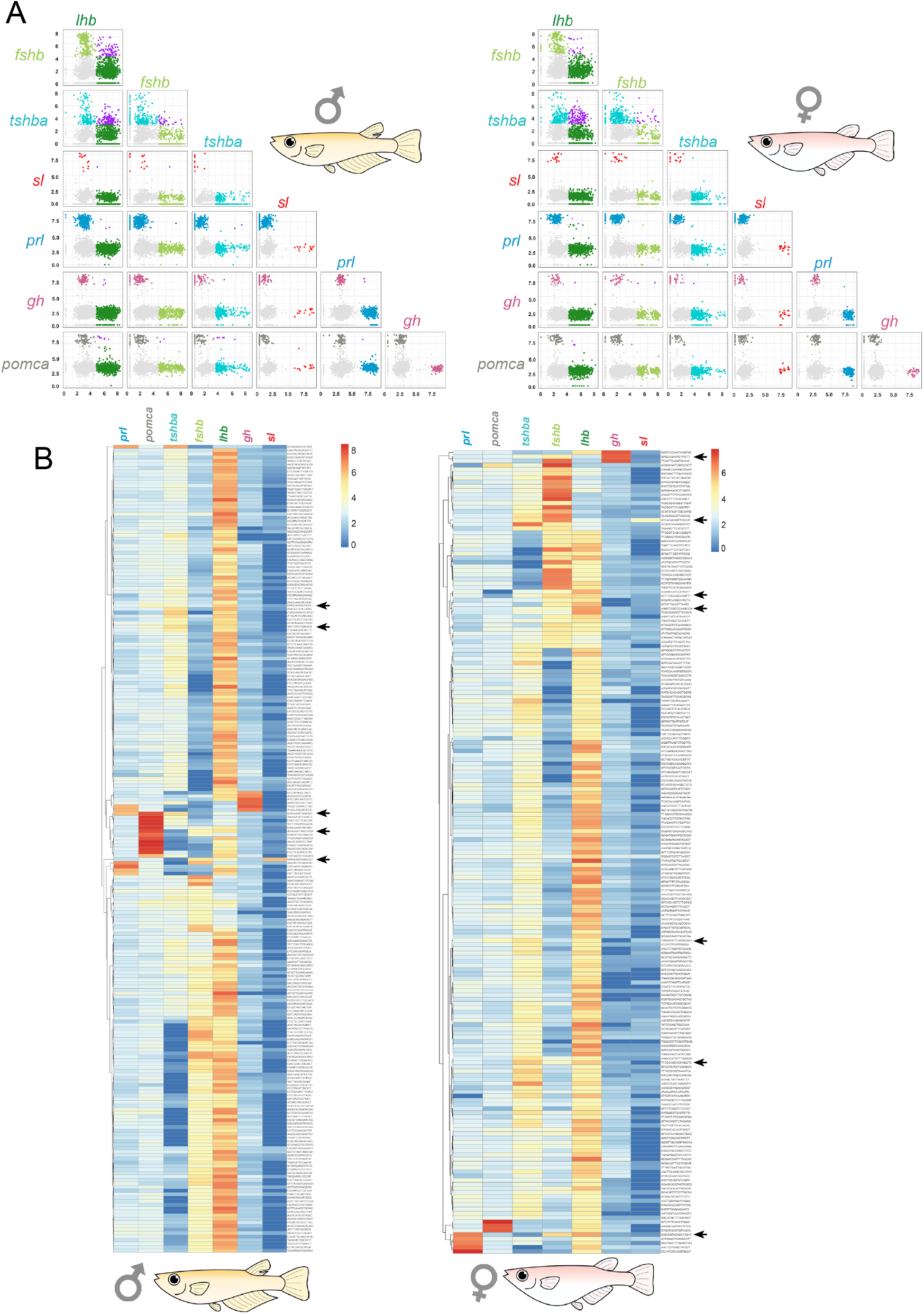
scRNA-seq data reveal the presence of bi-hormonal and multi-hormonal cells in the medaka pituitary. Pair wise plots of 2228 cells in female and 3245 cells in male pituitary (A). Colored by filtered cells (gray), *lhb* expressing cells (green), *fshb* expressing cells (darkolivegreen), *tshba* expressing cells (cyan), *sl* expressing cells (red), *prl* expressing cells (blue), *gh* expressing cells (magenta), *pomca* expressing cells (darkgrey) and cells expressing more than one endocrine gene (purple). Light grey cells represent the cells where gene expression for the investigated hormone is considered as part of the background. Axes are log normalized. Heatmap of seven hormone-encoding genes of the female and male pituitary (B). Each row represents one cell, and left side indicates cell clustering. Low expressions are shown in blue and high expressions are shown in red. Color bars on the left indicates how different cell types clustered on the basis of expression in each cell. Black arrows show cells that strongly express more than two hormone-encoding genes.

**Figure 6.**
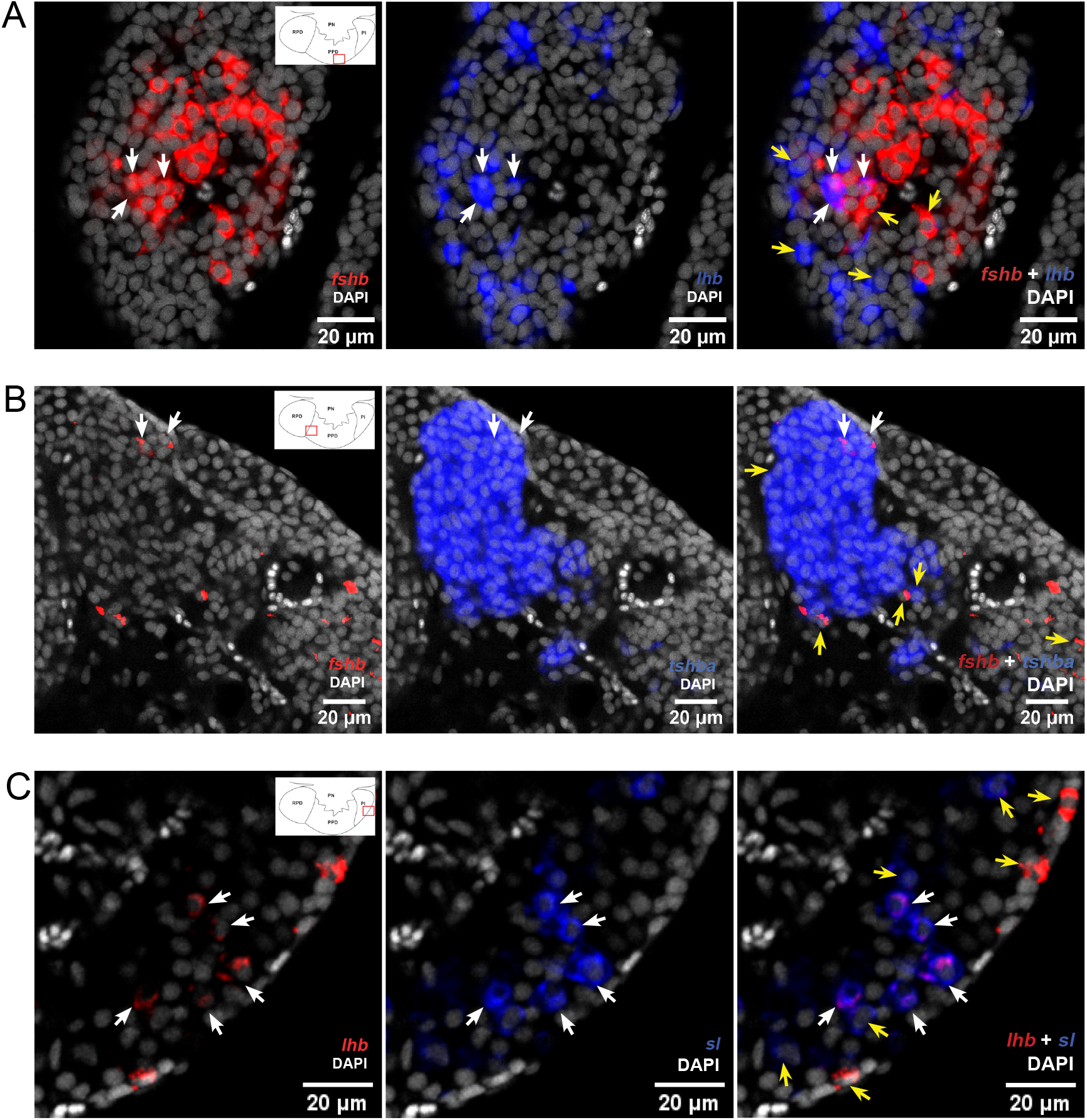
Multi-color FISH reveals some cells co-expressing more than one hormone-encoding genes in the medaka pituitary. Confocal plane confirming colocalization of *lhb* and *fshb* labelings (A), and *fshb* and *tshb* (B) labelings in both male and female adult medaka pituitary. Confocal plane showing the colocalization of *lhb* and *sl* in adult male (C). White arrows show cells whose mRNAs are co-expressed, while yellow arrows show cells that do not (can be used as control of probe’s specificity). The location of the bi-hormonal cells is in the proximity of red rectangle as illustrated in the schematic drawing of pituitary in left panels.

Several cells also co-express more than two hormone-encoding genes, although these are rare compared with bi-hormonal cells (Fig. 5B), and in this study they could not be confirmed using multi-color FISH.

## Discussion

### 3D spatial distribution of endocrine cell populations and blood vessels

We have recently used scRNA-seq to identify and characterize seven endocrine cell types in the teleost model organism medaka (31). Although a 3D atlas of the pituitary gland development has been previously described in zebrafish (38), the present atlas is the first 3D atlas of all pituitary endocrine cell populations in a teleost fish. It provides more precise and detailed information on the distribution and organization of the different cell types, and clearly demonstrate that endocrine cells are distributed differently when comparing mid-sagittal with para-sagittal sections (Fig. 7).

**Figure 7.**
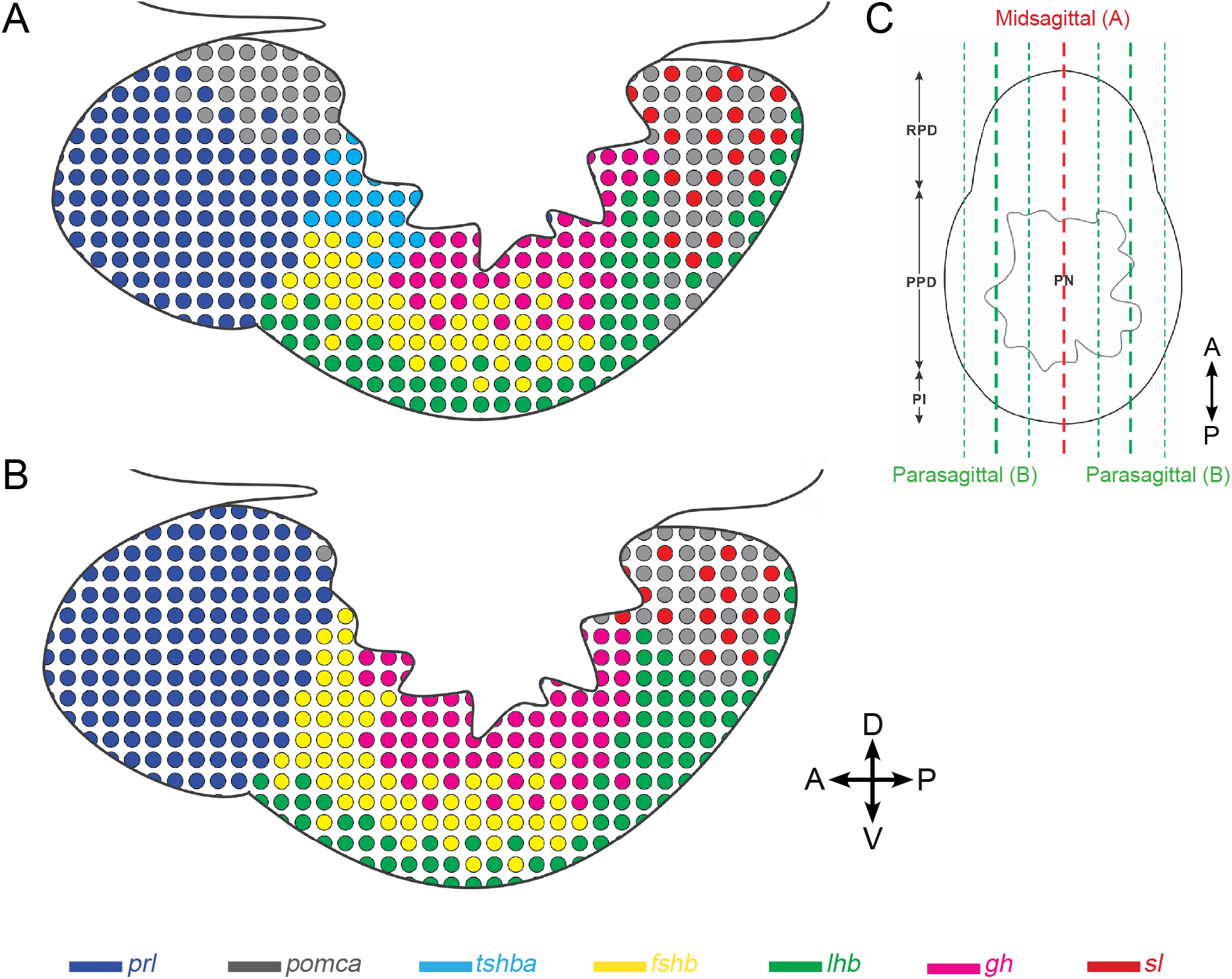
Schematic illustration showing differences in distribution of endocrine cell population between mid-sagittal and para-sagittal section of the medaka pituitary. The schemas are drawn based on mid- (A) and para-sagittal (B) point of view. C) Transverse view of medaka pituitary showing approximate location of mid-sagittal (red) and para-sagittal sections (green). Four direction arrows display the direction of the pituitary (A: anterior; P: posterior; D: dorsal; V: ventral).

As reported in coho (39) and chum salmon (40, 41), seabass (42), gilt-head seabream (43), common barbel (44) and striped bass (45), we observed some lactotropes in the ventro-peripheral area of the PPD. However, these cells were found only in some fish and not always at the same location, which makes them difficult to map. We found Lh gonadotropes in the PI, in addition to in the PPD, but only in adults, and we did not observe Lh cells in RPD as reported by some studies (16, 17, 46–48). The extra-PPD localization of Lh gonadotropes might be due to PPD extension (20) or Lh cell migration to other zones during the ontogeny of the adenohypophysis (49). Meanwhile, several studies reported somatotropes in the RPD (45, 50) and PI (51, 52), and somatolactotropes and melanotropes in the PPD (13, 16–18, 51). However, we did not observe this in medaka. The wide localization of Gh, Sl, and α-Msh cells in these studies might also be explained by antigenic similarities as suggested in (13, 16, 53).

The blood vasculature is ubiquitously spread over almost the entire adenohypophysis in medaka, without obvious differences between sexes and stages. This agrees with previous studies in zebrafish (54, 55) showing a highly vascularized pituitary. Such complex vasculature is, of course, central to the endocrinological function of the pituitary, as it allows for the efficient transport of secreted hormones to peripheral organs. In addition, it may facilitate intra-pituitary paracrine signaling (27).

### Sexual dimorphism of *tshba*- and *pomca*-expressing cell populations

While it seems impossible to show differences of endocrine cell population between sexes and stages when the data only rely on para-sagittal sections of the pituitary, whole pituitary labeling methods allow their identification. For instance, we show for the first time that *tshba*- and *pomca*-expressing cell populations are more numerous in adult females and males, respectively. Our qPCR data on *tshba* and *pomca* levels agree with previous studies in medaka (56) and further support the sexual dimorphism. In the previous medaka study, androgens were suggested to be involved in *tshb* suppression in adult males via activation of *tph1* transcription expressed in *pomc*-expressing cells. This is important for serotonin synthesis, which plays a role in repressing the expression of some endocrine cell markers, including *tshb*. We also found significantly lower *tshba* levels in adult compared to in juvenile males where the androgen levels are generally lower (for review, see (57, 58)), supporting the inhibitory role of androgens on *tshba* levels. It will be interesting to challenge this hypothesis in future research using orchidectomy which allows for androgen clearance in the medaka (59). However, although higher *tshb* levels are also observed in female half-barred wrasse (60) and *tshb* levels are higher in juvenile than in adult male Atlantic salmon (61), zebrafish shows no sexual dimorphism of *tshb* and *pomc* (62), suggesting species differences.

We also observed sexual dimorphism of *fshb* and *lhb* mRNA levels in adults, in agreement with a previous medaka study (56). This suggests a difference in gonadotrope cell activity as we did not observe a difference in population size, also supported by the absence of significant differences in Lh or Fsh cell numbers in previous studies (9, 63). In contrast, we observed an increase of *lhb* mRNA levels between juvenile and adult stages, which might mainly be due to an increase in cell numbers as shown in (63). Surprisingly, we did not observe a significant difference in *fshb* mRNA levels between juveniles and adults, while an increase in Fsh cell numbers has also been reported during sexual maturation (9). While neither sexual dimorphism nor stage differences were observed in *prl* or *gh* levels, we found stage difference in *sl* levels, with higher levels in juveniles than in adults. Somatolactin has been associated with sexual maturation in some teleosts, such as coho salmon (64), Nile tilapia (65) and flathead grey mullet (66). The upregulation of *sl* levels in teleosts is thought to be related to gonadal growth, as it is highly expressed at the onset of gonadal growth and lowly expressed post-ovulation (67, 68). This implies that in the current study, the adult fish used might be in a post-ovulation phase while the juveniles appear to enter gonadal development.

### scRNA-seq and multi-color FISH reveal the presence of multi-hormonal cells in the adult medaka pituitary

The presence of cells expressing more than one hormone in the anterior pituitary has been shown in many studies, both in teleosts and in mammals (for review see (28, 69–71)). Using scRNA-seq technology, multi-hormonal cells have for instance been described in the mouse pituitary (72). However, using similar approaches, pituitary multi-hormonal cells have not been reported in teleosts (31, 73), although some studies have documented cells producing more than one hormone in the teleost pituitary (8–11).

Here, we show the presence of gonadotrope cells expressing both *lhb* and *fshb* which has previously been reported in medaka (9) and in other teleost species (8, 10, 11). One proposed underlying mechanism that could explain the existence of such bi-hormonal gonadotropes is their capacity to change phenotype, with a transitory bi-hormonal state as suggested by *in vitro* observation where Lh cells were found to become Fsh (9). We also found cells co-expressing *fshb-tshba* and *sl-lhb*. Although previous immunohistochemtry studies on the pituitary of several teleost species showed cross-reaction between Tsh and Lh/Fsh (43, 53) and between Lh and Sl antibodies (16, 53, 74, 75), we show specific labeling of *tshb, fshb, lhb*, and *sl* in the current study confirming the specificity of the probes. Therefore, the observation of colocalization of *fshb-tshba* and *sl-lhb* supports that these bi-hormonal cells exist in the medaka pituitary. Meanwhile, a number of studies have shown co-staining between Prl, Gh and Sl (13, 20, 22, 40, 51). However, our scRNA-seq data exhibit only a few cells co-expressing *prl* and *gh*. While antigenic similarities could be the reason of co-staining between these cell types (13, 16, 53), their low occurrence in the medaka pituitary might be the reason of being unable to observe them with FISH.

Within the *pomca*-expressing cell population, we observed Acth staining alone at the border of the RPD/PPD and staining of both Acth and α-Msh in the PI. Co-staining between Acth and α-Msh is not uncommon as Acth-immunoreactive cells have been found in RPD and PI areas in other teleost species (8, 18, 20–22, 42, 43, 76). However, it must be noted that the target antigen of anti-Acth used in this study and most previous studies contains the target antigen for anti-α-Msh (https://www.uniprot.org/uniprot/P01189#PRO_0000024970). This might explain why we found both Acth and α-Msh cells in the PI. A previous study pre-incubating anti-Acth with α-Msh antigen demonstrated that Acth cells are localized in the RPD while α-Msh cells are found in the PI (21). This might also be the case in medaka.

We also show that a few cells in the adult medaka pituitary express more than two hormone-encoding genes. Despite having been reported in mammals (72), the current study is the first to show the presence of such multi-hormonal cells in the teleost pituitary. Although we could not confirm their existence using FISH, most likely because of the low number of such cells present in the pituitary, the demonstration of their existence using scRNA-seq raises questions about their origin and roles in the medaka pituitary.

Finally, the 3D atlas platform that is provided online will help research community to take a close look on the spatial distribution of endocrine cells and the vascularization of blood vessels in the pituitary.

## Author Contributions

RF and FAW conceptualized and planned the work. RF, MAP, JB and FAW obtained funding. NR did all cloning. MRR and RF performed the experiments and acquired the imaging data. MRR and GC processed the imaging data and developed the online 3D model, supervised by MAP and JB. CH and KS analyzed the single cell transcriptome data. MRR, CH and RF wrote the paper with the inputs from all authors.

## Acknowledgments

The authors thank Ms Lourdes Carreon G. Tan for her assistance in the fish husbandry.

---

**Supp. Fig. 1.** Density plots of hormone-encoding genes in the pituitary of adult male (A) and female (B) medaka. Red line represents a cut-off to differentiate between the cells with high gene expression (considered as endocrine cells) and low gene expression (considered as background and non-endocrine cells).

**Supp. Fig. 2-8.** Orthogonal view of each endocrine cell types in juvenile and adult male and female medaka. (2, *prl*; 3, *pomca;* 4, *tshba;* 5, *fshb;* 6, *lhb*; 7, *gh*; 8, *sl*). The pictures were captured from different perspectives: transverse (i), frontal (ii) and para-sagittal (iii). Up down arrow symbol shows the direction of the pituitary (A: anterior; P: posterior; D: dorsal; V: ventral).

**Supp. Fig. 9-13.** Transverse views of multi-color FISH of endocrine cells in juvenile and adult male and female medaka. (9, *prl*-*tshba*-*pomca*; 10, *sl*-*lhb*-*pomca*; 11, *fshb*-*tshba*; 12, *fshb-lhb;* 13, *fshb-gh*) (A: anterior; P: posterior; D: dorsal; V: ventral).

**Supp. Fig. 14.** The combination of FISH for *pomca* and IF for Acth or α-Msh allows the distinction of two clear *pomca* expressing cell populations. The distinction of Acth (green) and α-Msh (blue) producing cells from *pomca*-labelled (red) in the pituitary from juvenile male and female medaka. Dashed line represents the pituitary as shown in the right panel. Four direction arrows display the direction of the pituitary (A: anterior; P: posterior; D: dorsal; V: ventral).

## References

1. Yeung C-M, Chan C-B, Leung P-S, Cheng CHK. Cells of the anterior pituitary. The International Journal of Biochemistry & Cell Biology. 2006;38(9):1441–9.

2. Ooi GT, Tawadros N, Escalona RM. Pituitary cell lines and their endocrine applications. Molecular and Cellular Endocrinology. 2004;228(1):1–21.

3. Melmed S. The pituitary: Academic press; 2010.

4. Brinkmeier ML, Davis SW, Carninci P, MacDonald JW, Kawai J, Ghosh D, et al. Discovery of transcriptional regulators and signaling pathways in the developing pituitary gland by bioinformatic and genomic approaches. Genomics. 2009;93(5):449–60.

5. Le Tissier PR, Hodson DJ, Lafont C, Fontanaud P, Schaeffer M, Mollard P. Anterior pituitary cell networks. Frontiers in Neuroendocrinology. 2012;33(3):252–66.

6. Kaneko T. Cell Biology of Somatolactin. In: Jeon KW, editor. International Review of Cytology. 169: Academic Press; 1996. p. 1–24.

7. Weltzien F-A, Hildahl J, Hodne K, Okubo K, Haug TM. Embryonic development of gonadotrope cells and gonadotropic hormones – Lessons from model fish. Molecular and Cellular Endocrinology. 2014;385(1):18–27.

8. Hernández MPGa, García Ayala A, Zandbergen MA, Agulleiro B. Investigation into the duality of gonadotropic cells of Mediterranean yellowtail (Seriola dumerilii, Risso 1810): immunocytochemical and ultrastructural studies. General and Comparative Endocrinology. 2002;128(1):25–35.

9. Fontaine R, Ager-Wick E, Hodne K, Weltzien F-A. Plasticity in medaka gonadotropes via cell proliferation and phenotypic conversion. Journal of Endocrinology. 2020;245(1):21.

10. Candelma M, Fontaine R, Colella S, Santojanni A, Weltzien F-A, Carnevali O. Gonadotropin characterization, localization and expression in the European hake (Merluccius merluccius). Reproduction. 2017;153(2):123.

11. Golan M, Biran J, Levavi-Sivan B. A Novel Model for Development, Organization, and Function of Gonadotropes in Fish Pituitary. Frontiers in Endocrinology. 2014;5(182).

12. Schreibman MP, Leatherland JF, McKeown BA. Functional Morphology of the Teleost Pituitary Gland. American Zoologist. 2015;13(3):719–42.

13. Honji RM, Nóbrega RH, Pandolfi M, Shimizu A, Borella MI, Moreira RG. Immunohistochemical study of pituitary cells in wild and captive Salminus hilarii (Characiformes: Characidae) females during the annual reproductive cycle. SpringerPlus. 2013;2(1):460.

14. Aoki K, Umeura H. Cell Types in the Pituitary of the Medaka (Oryzias latipes). Endocrinologia Japonica. 1970;17(1):45–55.

15. Mukai T, Oota Y. Histological Changes in the Pituitary, Thyroid Gland and Gonads of the Fourspine Sculpin (Cottus kazika) during Downstream Migration. Zoological Science. 1995;12(1):91–7.

16. Camacho LR, Pozzi AG, de Freitas EG, Shimizu A, Pandolfi M. Morphological and immunohistochemical comparison of the pituitary gland between a tropical Paracheirodon axelrodi and a subtropical Aphyocharax anisitsi characids (Characiformes: Characidae) %J Neotropical Ichthyology. 2020;18.

17. Sánchez Cala F, Portillo A, Martín del Río MP, Mancera JM. Immunocytochemical characterization of adenohypophyseal cells in the greater weever fish (Trachinus draco). Tissue and Cell. 2003;35(3):169–78.

18. Segura-Noguera MM, Laíz-Carrión R, Martín del Río MP, Mancera JM. An Immunocytochemical Study of the Pituitary Gland of the White Seabream (Diplodus Sargus). The Histochemical Journal. 2000;32(12):733–42.

19. Pandolfi M, Cánepa MM, Meijide FJ, Alonso F, Vázquez GR, Maggese MC, et al. Studies on the reproductive and developmental biology of Cichlasoma dimerus (Percifomes, Cichlidae). Biocell. 2009;33(1):1–18.

20. Weltzien F-A, Norberg B, Helvik JV, Andersen Ø, Swanson P, Andersson E. Identification and localization of eight distinct hormone-producing cell types in the pituitary of male Atlantic halibut (Hippoglossus hippoglossus L.). Comparative Biochemistry and Physiology Part A: Molecular & Integrative Physiology. 2003;134(2):315–27.

21. Kasper RS, Shved N, Takahashi A, Reinecke M, Eppler E. A systematic immunohistochemical survey of the distribution patterns of GH, prolactin, somatolactin, β–TSH, β–FSH, β–LH, ACTH, and α–MSH in the adenohypophysis of Oreochromis niloticus, the Nile tilapia. Cell and Tissue Research. 2006;325(2):303–13.

22. Parhar IS, Nagahama Y, Grau EG, Ross RM. Immunocytochemical and Ultrastructural Identification of Pituitary Cell Types in the Protogynous <span class=“genus-species”>Thalassoma duperrey</span> during Adult Sexual Ontogeny. 1998;15 %J Zoological Science(2):263–76, 14.

23. Childs GV. Development of gonadotropes may involve cyclic transdifferentiation of growth hormone cells. Archives of physiology and biochemistry. 2002;110(1-2):42–9.

24. Childs GV. Multipotential pituitary cells that contain adrenocorticotropin (ACTH) and other pituitary hormones. Trends in Endocrinology & Metabolism. 1991;2(3):112–7.

25. Frawley LS, Boockfor FR. Mammosomatotropes: Presence and Functions in Normal and Neoplastic Pituitary Tissue. Endocrine Reviews. 1991;12(4):337–55.

26. Fukami K, Tasaka K, Mizuki J, Kasahara K, Masumoto N, Miyake A, et al. Bihormonal Cells Secreting Both Prolactin and Gonadotropins in Normal Rat Pituitary Cells. Endocrine Journal. 1997;44(6):819–26.

27. Ben-Shlomo A, Melmed S. Chapter 2 - Hypothalamic Regulation of Anterior Pituitary Function. In: Melmed S, editor. The Pituitary (Third Edition). San Diego: Academic Press; 2011. p. 21–45.

28. Fontaine R, Ciani E, Haug TM, Hodne K, Ager-Wick E, Baker DM, et al. Gonadotrope plasticity at cellular, population and structural levels: A comparison between fishes and mammals. General and Comparative Endocrinology. 2020;287:113344.

29. Wittbrodt J, Shima A, Schartl M. Medaka — a model organism from the far east. Nature Reviews Genetics. 2002;3(1):53–64.

30. Naruse K. Genetics, Genomics, and Biological Resources in the Medaka, Oryzias latipes. In: Naruse K, Tanaka M, Takeda H, editors. Medaka: A Model for Organogenesis, Human Disease, and Evolution. Tokyo: Springer Japan; 2011. p. 19–37.

31. Siddique K, Ager-Wick E, Fontaine R, Weltzien F-A, Henkel CV. Characterization of hormone-producing cell types in the medaka pituitary gland using single-cell RNA-seq. bioRxiv. 2020.

32. Kenji Murata, Masato Kinoshita, Kiyoshi Naruse, Minoru Tanaka, Kamei Y. Looking at Adult Medaka. In: Kenji Murata, Masato Kinoshita, Kiyoshi Naruse, Minoru Tanaka, Kamei Y, editors. Medaka: Biology, Management, and Experimental Protocols. 2: John Wiley & Sons; 2019. p. 49–95.

33. Burow S, Fontaine R, von Krogh K, Mayer I, Nourizadeh-Lillabadi R, Hollander-Cohen L, et al. Medaka follicle-stimulating hormone (Fsh) and luteinizing hormone (Lh): Developmental profiles of pituitary protein and gene expression levels. General and Comparative Endocrinology. 2019;272:93–108.

34. Fontaine R, Affaticati P, Yamamoto K, Jolly C, Bureau C, Baloche S, et al. Dopamine Inhibits Reproduction in Female Zebrafish (Danio rerio) via Three Pituitary D2 Receptor Subtypes. Endocrinology. 2013;154(2):807–18.

35. Fontaine R, Weltzien F-A. Labeling of Blood Vessels in the Teleost Brain and Pituitary Using Cardiac Perfusion with a DiI-fixative. Journal of Visualized Experiments. 2019 (148):e59768.

36. Bjerke IE, Øvsthus M, Papp EA, Yates SC, Silvestri L, Fiorilli J, et al. Data integration through brain atlasing: Human Brain Project tools and strategies. European Psychiatry. 2018;50:70–6.

37. Yates SC, Groeneboom NE, Coello C, Lichtenthaler SF, Kuhn P-H, Demuth H-U, et al. QUINT: Workflow for Quantification and Spatial Analysis of Features in Histological Images From Rodent Brain. Frontiers in Neuroinformatics. 2019;13(75).

38. Chapman SC, Sawitzke AL, Campbell DS, Schoenwolf GC. A three-dimensional atlas of pituitary gland development in the zebrafish. Journal of Comparative Neurology. 2005;487(4):428–40.

39. Farbridge KJ, McDonald-Jones G, McLean CL, Lowry PJ, Etches RJ, Leatherland JF. The development of monoclonal antibodies against salmon (Oncorhynchus kisutch and O. keta) pituitary hormones and their immunohistochemical identification. General and Comparative Endocrinology. 1990;79(3):361–74.

40. Naito N, Takahashi A, Nakai Y, Kawauchi H, Hirano T. Immunocytochemical identification of the prolactin-secreting cells in the teleost pituitary with an antiserum to chum salmon prolactin. General and Comparative Endocrinology. 1983;50(2):282–91.

41. Wagner GF, McKeown BA. The immunocytochemical localization of pituitary somatotrops in the genus Oncorhynchus using an antiserum to growth hormone of chum salmon (Oncorhynchus keta). Cell Tissue Res. 1983;231(3):693–7.

42. Cambré ML, Verdonck W, Ollevier F, Vandesande F, Batten TFC, Kühn ER. Immunocytochemical identification and localization of the different cell types in the pituitary of the seabass (Dicentrarchus labrax). General and Comparative Endocrinology. 1986;61(3):368–75.

43. Quesada J, Lozano MT, Ortega A, Agulleiro B. Immunocytochemical and ultrastructural characterization of the cell types in the adenohypophysis of Sparus aurata L. (Teleost). General and Comparative Endocrinology. 1988;72(2):209–25.

44. Toubeau G, Poilve A, Baras E, Nonclercq D, De Moor S, Beckers JF, et al. Immunocytochemical study of cell type distribution in the pituitary of Barbus barbus (Teleostei, Cyprinidae). General and Comparative Endocrinology. 1991;83(1):35–47.

45. Huang L, Specker JL. Growth Hormone- and Prolactin-Producing Cells in the Pituitary Gland of Striped Bass (Morone saxatilis): Immunocytochemical Characterization at Different Life Stages. General and Comparative Endocrinology. 1994;94(2):225–36.

46. Borella MI, Venturieri R, Mancera JM. Immunocytochemical identification of adenohypophyseal cells in the pirarucu (Arapaima gigas), an Amazonian basal teleost. Fish Physiology and Biochemistry. 2009;35(1):3–16.

47. Olivereau M, Nagahama Y. Immunocytochemistry of gonadotropic cells in the pituitary of some teleost species. General and Comparative Endocrinology. 1983;50(2):252–60.

48. Dubourg P, Burzawa-Gerard E, Chambolle P, Kah O. Light and electron microscopic identification of gonadotrophic cells in the pituitary gland of the goldfish by means of immunocytochemistry. General and Comparative Endocrinology. 1985;59(3):472–81.

49. Nozaki M, Naito N, Swanson P, Miyata K, Nakai Y, Oota Y, et al. Salmonid pituitary gonadotrophs I. Distinct cellular distributions of two gonadotropins, GTH I and GTH II. General and Comparative Endocrinology. 1990;77(3):348–57.

50. Grandi G, Colombo G, Chicca M. Immunocytochemical studies on the pituitary gland of Anguilla anguilla L., in relation to early growth stages and diet-induced sex differentiation. General and Comparative Endocrinology. 2003;131(1):66–76.

51. García-Hernández MP, García-Ayala A, Elbal MT, Agulleiro B. The adenohypophysis of Mediterranean yellowtail, Seriola dumerilii (Risso, 1810): an immunocytochemical study. Tissue and Cell. 1996;28(5):577–85.

52. Grandi G, Chicca M. Early development of the pituitary gland in Acipenser naccarii (Chondrostei, Acipenseriformes): an immunocytochemical study. Anatomy and Embryology. 2004;208(4):311–21.

53. Batten TFC. Immunocytochemical demonstration of pituitary cell types in the teleost Poecilia latipinna, by light and electron microscopy. General and Comparative Endocrinology. 1986;63(1):139–54.

54. Gutnick A, Blechman J, Kaslin J, Herwig L, Belting H-G, Affolter M, et al. The Hypothalamic Neuropeptide Oxytocin Is Required for Formation of the Neurovascular Interface of the Pituitary. Developmental Cell. 2011;21(4):642–54.

55. Golan M, Zelinger E, Zohar Y, Levavi-Sivan B. Architecture of GnRH-Gonadotrope-Vasculature Reveals a Dual Mode of Gonadotropin Regulation in Fish. Endocrinology. 2015;156(11):4163–73.

56. Kawabata-Sakata Y, Nishiike Y, Fleming T, Kikuchi Y, Okubo K. Androgen-dependent sexual dimorphism in pituitary tryptophan hydroxylase expression: relevance to sex differences in pituitary hormones. Proceedings of the Royal Society B: Biological Sciences. 2020;287(1928):20200713.

57. Taranger GL, Carrillo M, Schulz RW, Fontaine P, Zanuy S, Felip A, et al. Control of puberty in farmed fish. General and Comparative Endocrinology. 2010;165(3):483–515.

58. Borg B. Androgens in teleost fishes. Comparative Biochemistry and Physiology Part C: Pharmacology, Toxicology and Endocrinology. 1994;109(3):219–45.

59. Royan MR, Kanda S, Kayo D, Song W, Ge W, Weltzien F-A, et al. Gonadectomy and Blood Sampling Procedures in the Small Size Teleost Model Japanese Medaka (Oryzias latipes). JoVE. 2020(166):e62006.

60. Ohta K, Mine T, Yamaguchi A, Matsuyama M. Sexually dimorphic expression of pituitary glycoprotein hormones in a sex-changing fish (Pseudolabrus sieboldi). Journal of Experimental Zoology Part A: Ecological Genetics and Physiology. 2008;309A(9):534–41.

61. Martin SAM, Wallner W, Youngson AF, Smith T. Differential expression of Atlantic salmon thyrotropin β subunit mRNA and its cDNA sequence. Journal of Fish Biology. 1999;54(4):757–66.

62. He W, Dai X, Chen X, He J, Yin Z. Zebrafish pituitary gene expression before and after sexual maturation. Journal of Endocrinology. 2014;221(3):429.

63. Fontaine R, Ager-Wick E, Hodne K, Weltzien F-A. Plasticity of Lh cells caused by cell proliferation and recruitment of existing cells. Journal of Endocrinology. 2019;240(2):361.

64. Rand-Weaver M, Swanson P, Kawauchi H, Dickhoff WW. Somatolactin, a novel pituitary protein: purification and plasma levels during reproductive maturation of coho salmon. Journal of Endocrinology. 1992;133(3):393.

65. Mousa MA, Mousa SA. Immunocytochemical Study on the Localization and Distribution of the Somatolactin Cells in the Pituitary Gland and the Brain ofOreochromis niloticus(Teleostei, Cichlidae). General and Comparative Endocrinology. 1999;113(2):197–211.

66. Mousa MA, Mousa SA. Implication of somatolactin in the regulation of sexual maturation and spawning of Mugil cephalus. Journal of Experimental Zoology. 2000;287(1):62–73.

67. Benedet S, Björnsson BT, Taranger GL, Andersson E. Cloning of somatolactin alpha, beta forms and the somatolactin receptor in Atlantic salmon: seasonal expression profile in pituitary and ovary of maturing female broodstock. Reproductive biology and endocrinology. 2008;6:42-.

68. Onuma T, Kitahashi T, Taniyama S, Saito D, Ando H, Urano A. Changes in expression of genes encoding gonadotropin subunits and growth hormone/prolactin/somatolactin family hormones during final maturation and freshwater adaptation in prespawning chum salmon. Endocrine. 2003;20(1):23–33.

69. Fontaine R, Royan MR, von Krogh K, Weltzien F-A, Baker DM. Direct and Indirect Effects of Sex Steroids on Gonadotrope Cell Plasticity in the Teleost Fish Pituitary. Frontiers in Endocrinology. 2020;11(858).

70. Childs GV, MacNicol AM, MacNicol MC. Molecular Mechanisms of Pituitary Cell Plasticity. Frontiers in Endocrinology. 2020;11(656).

71. Rizzoti K. Adult pituitary progenitors/stem cells: from in vitro characterization to in vivo function. 2010;32(12):2053–62.

72. Ho Y, Hu P, Peel MT, Chen S, Camara PG, Epstein DJ, et al. Single-cell transcriptomic analysis of adult mouse pituitary reveals sexual dimorphism and physiologic demand-induced cellular plasticity. Protein & Cell. 2020;11(8):565–83.

73. Fabian P, Tseng KC, Smeeton J, Lancman JJ, Dong PDS, Cerny R, et al. Lineage analysis reveals an endodermal contribution to the vertebrate pituitary. Science. 2020;370(6515):463–7.

74. Batten T, Ball JN, Benjamin M. Ultrastructure of the adenohypophysis in the teleost Poecilia latipinna. Cell and Tissue Research. 1975;161(2):239–61.

75. Margolis-Kazan H, Peute J, Schreibman MP, Halpern LR. Ultrastructural localization of gonadotropin and luteinizing hormone releasing hormone in the pituitary gland of a teleost fish (the platyfish). Journal of Experimental Zoology. 1981;215(1):99–102.

76. Yan HY, Thomas P. Histochemical and immunocytochemical identification of the pituitary cell types in three sciaenid fishes: Atlantic croaker (Micropogonias undulatus), spotted seatrout (Cynoscion nebulosus), and red drum (Sciaenops ocellatus). General and Comparative Endocrinology. 1991;84(3):389–400.

