## supplemental figures and tables for "3D atlas of the pituitary gland of the model fish medaka"

A

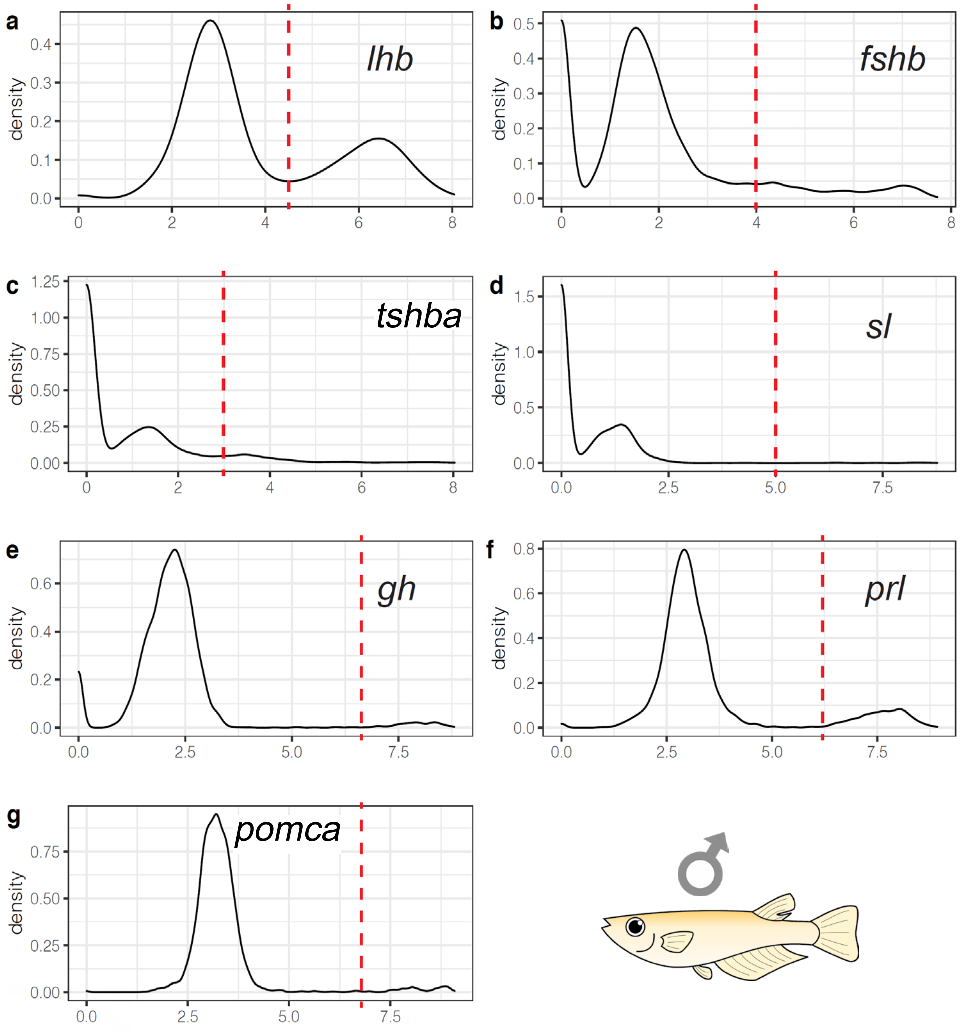

a
lhb
density
0.4
0.3
0.2
0.1
0.0
0
2
4
6
8
b
fshb
0.5
0.4
0.3
0.2
0.1
0.0
0
2
4
6
8
c
tshba
1.25
1.00
0.75
0.50
0.25
0.00
0
2
4
6
8
d
sl
1.5
1.0
0.5
0.0
0.0
2.5
5.0
7.5
e
gh
0.6
0.4
0.2
0.0
0.0
2.5
5.0
7.5
f
prl
0.8
0.6
0.4
0.2
0.0
0.0
2.5
5.0
7.5
g
pomca
0.75
0.50
0.25
0.00
0.0
2.5
5.0
7.5
♂


B

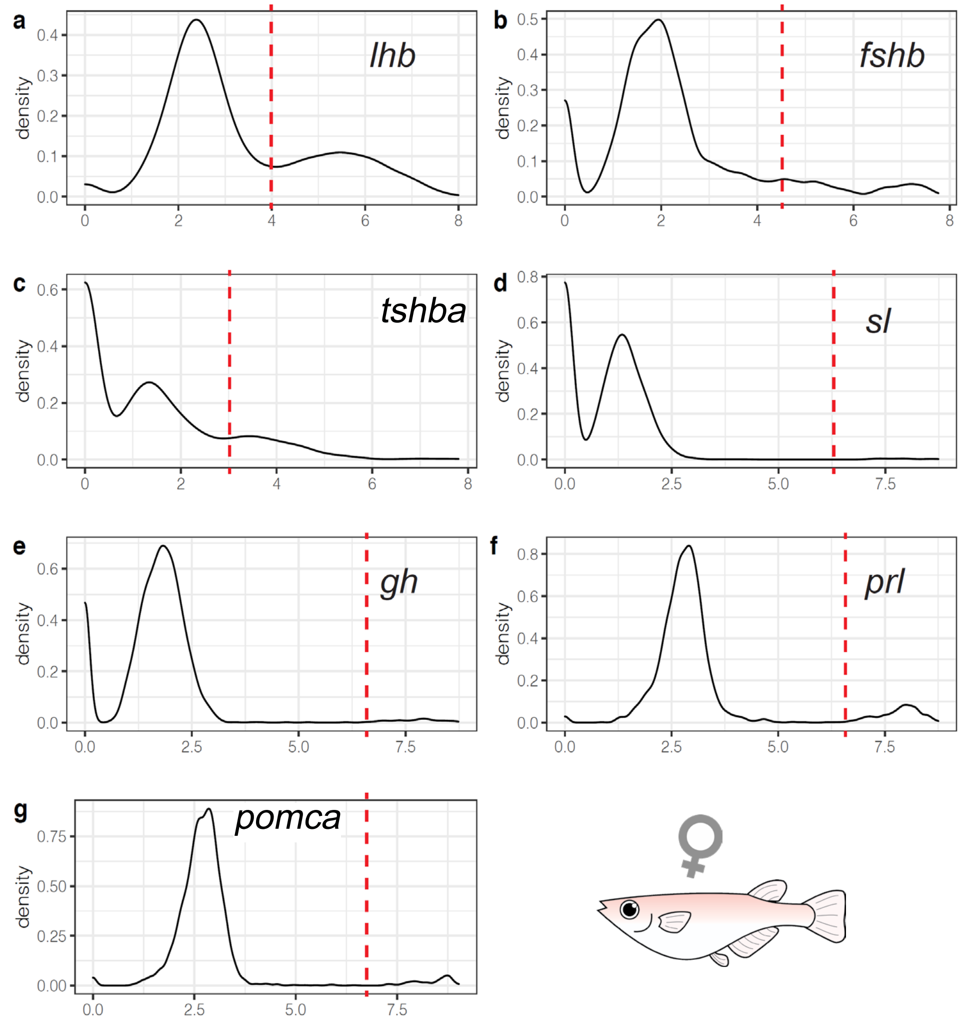

a
lhb
density
0.4
0.3
0.2
0.1
0.0
0
2
4
6
8
b
fshb
0.5
0.4
0.3
0.2
0.1
0.0
0
2
4
6
8
c
tshba
0.6
0.4
0.2
0.0
0
2
4
6
8
d
sl
0.8
0.6
0.4
0.2
0.0
0.0
2.5
5.0
7.5
e
gh
0.6
0.4
0.2
0.0
0.0
2.5
5.0
7.5
f
prl
0.8
0.6
0.4
0.2
0.0
0.0
2.5
5.0
7.5
g
pomca
0.75
0.50
0.25
0.00
0.0
2.5
5.0
7.5
♀


SUPP. FIG. 1

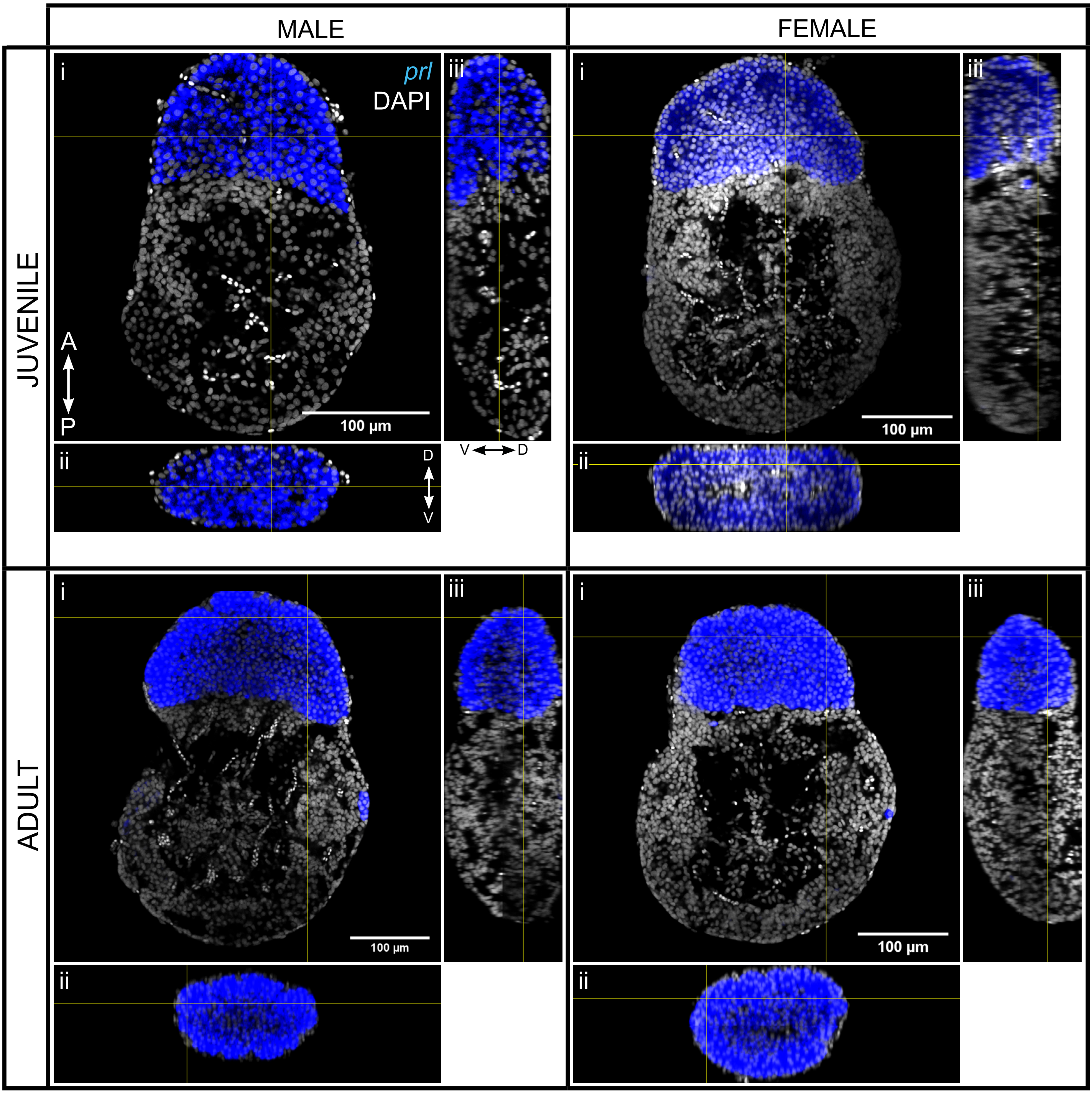
MALE
FEMALE
JUVENILE
ADULT
i
ii
iii
prl
DAPI
A
P
D
V
V
D
100 µm
100 µm
100 µm
100 µm

SUPP. FIG. 2

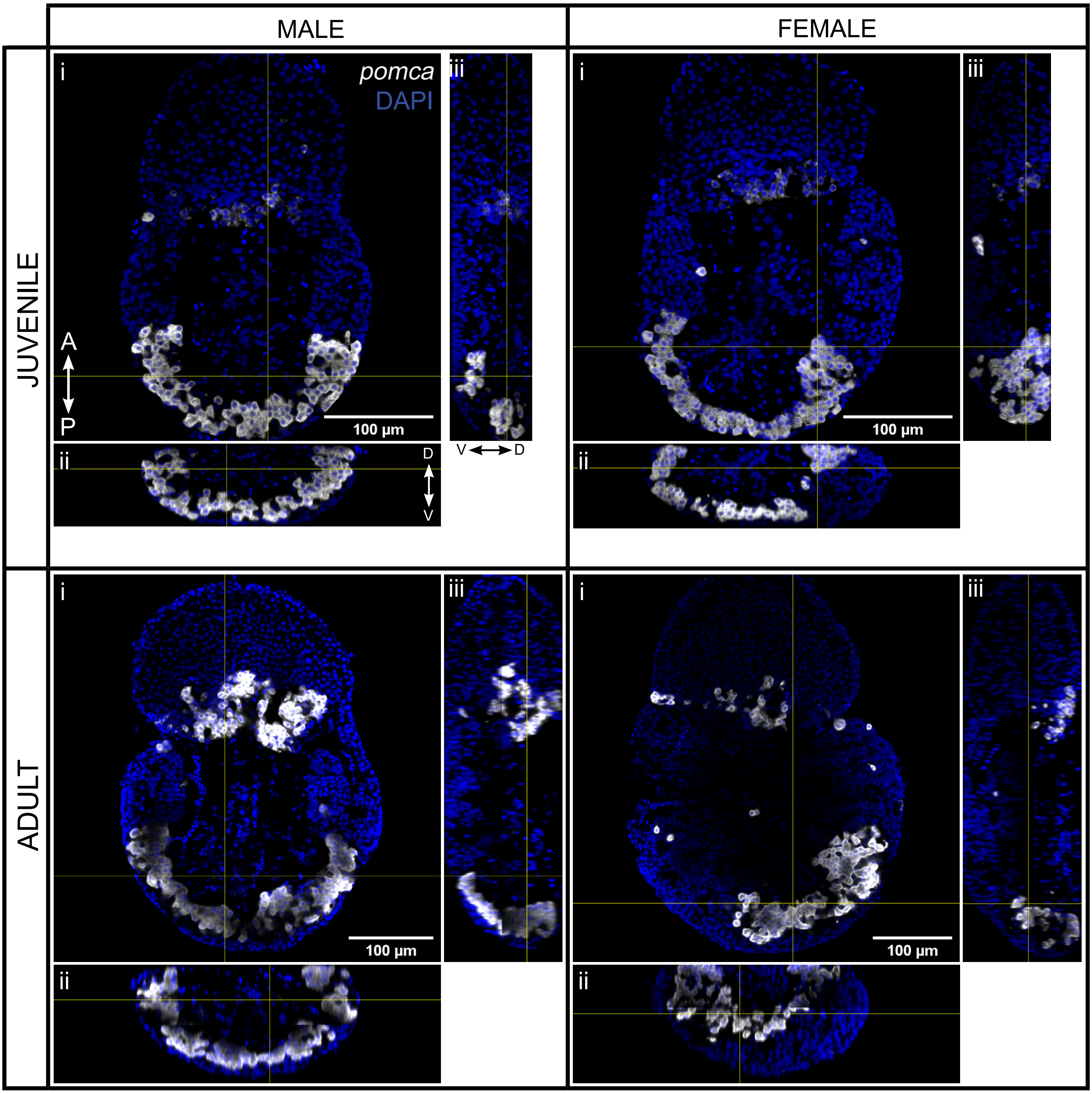


SUPP. FIG. 3

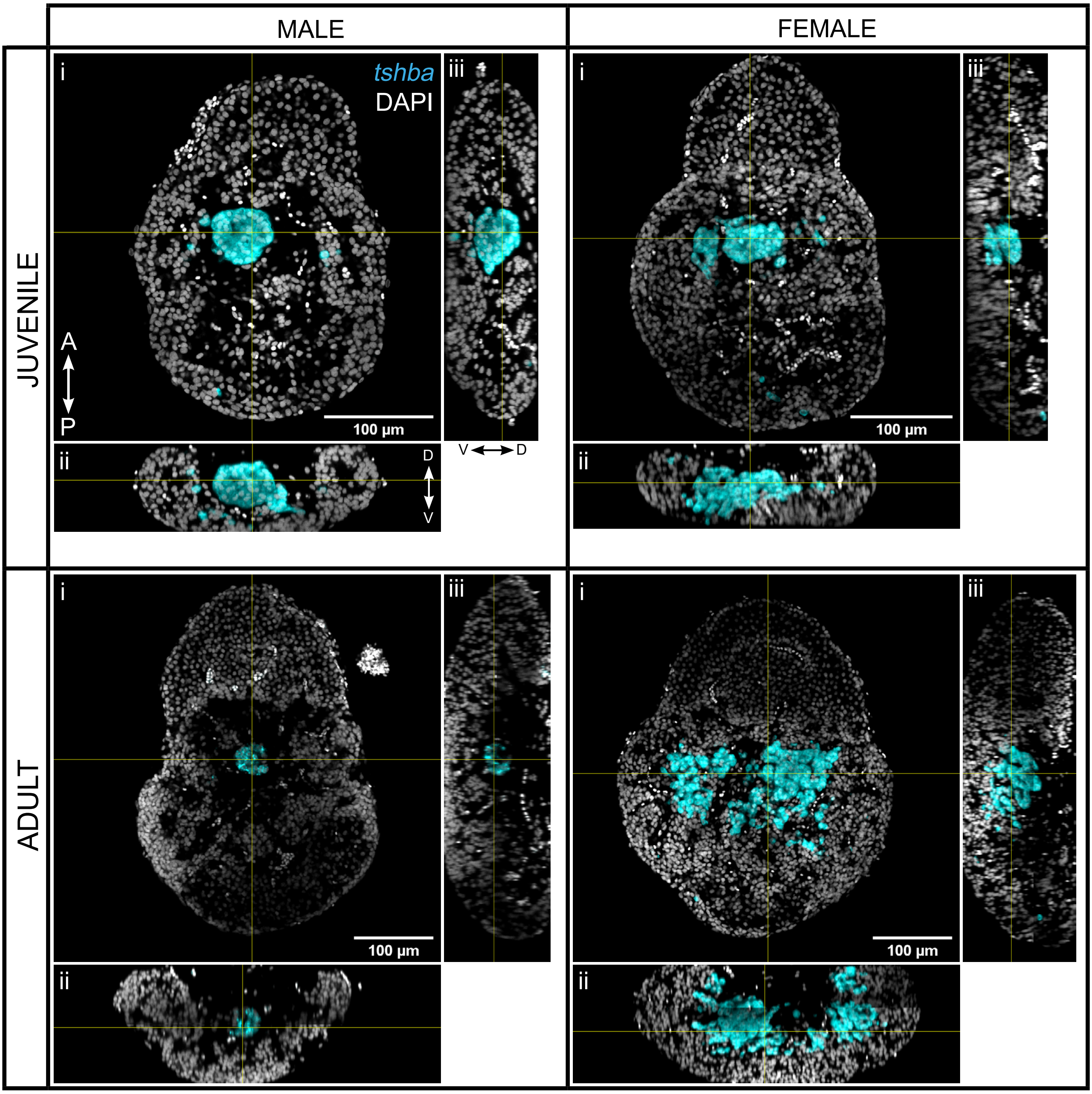
MALE
FEMALE
JUVENILE
ADULT
i
ii
iii
tshba
DAPI
A
P
D
V
V
D
100 µm
100 µm
100 µm
100 µm

SUPP. FIG. 4

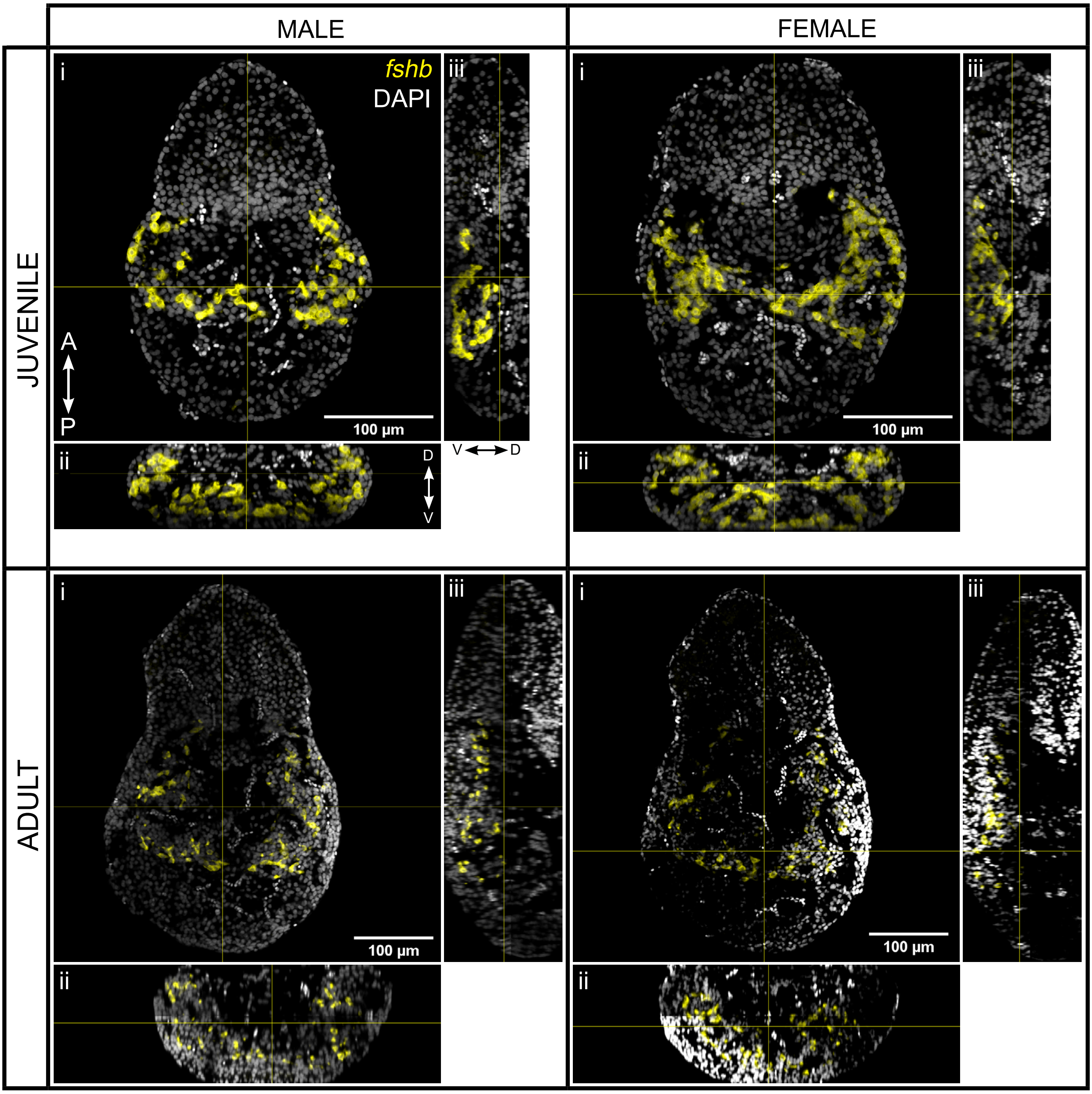

MALE
FEMALE
JUVENILE
ADULT
i
ii
iii
fshb
DAPI
A
P
D
V
V
D
100 µm
100 µm
100 µm
100 µm


SUPP. FIG. 5

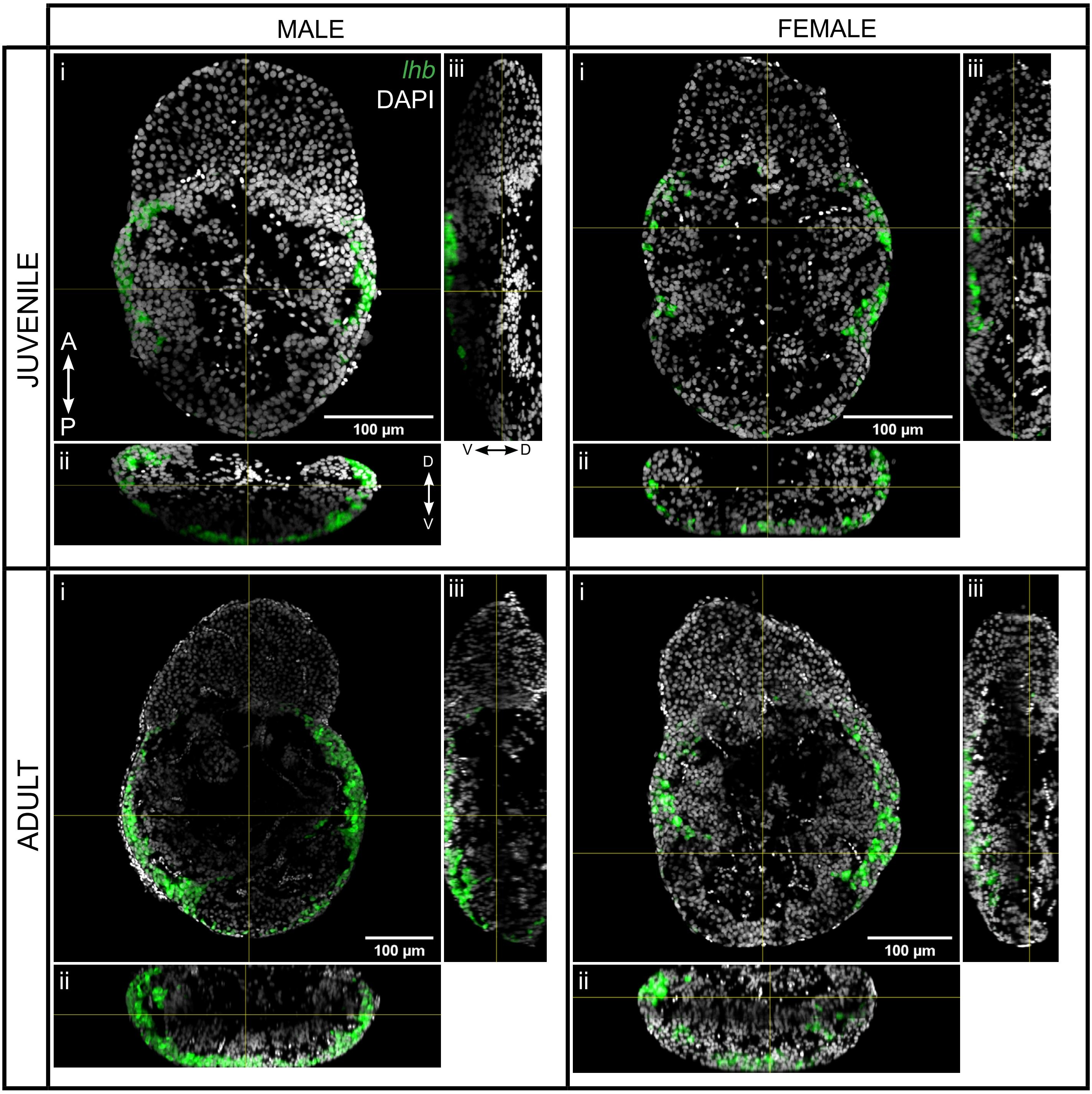


SUPP. FIG. 6

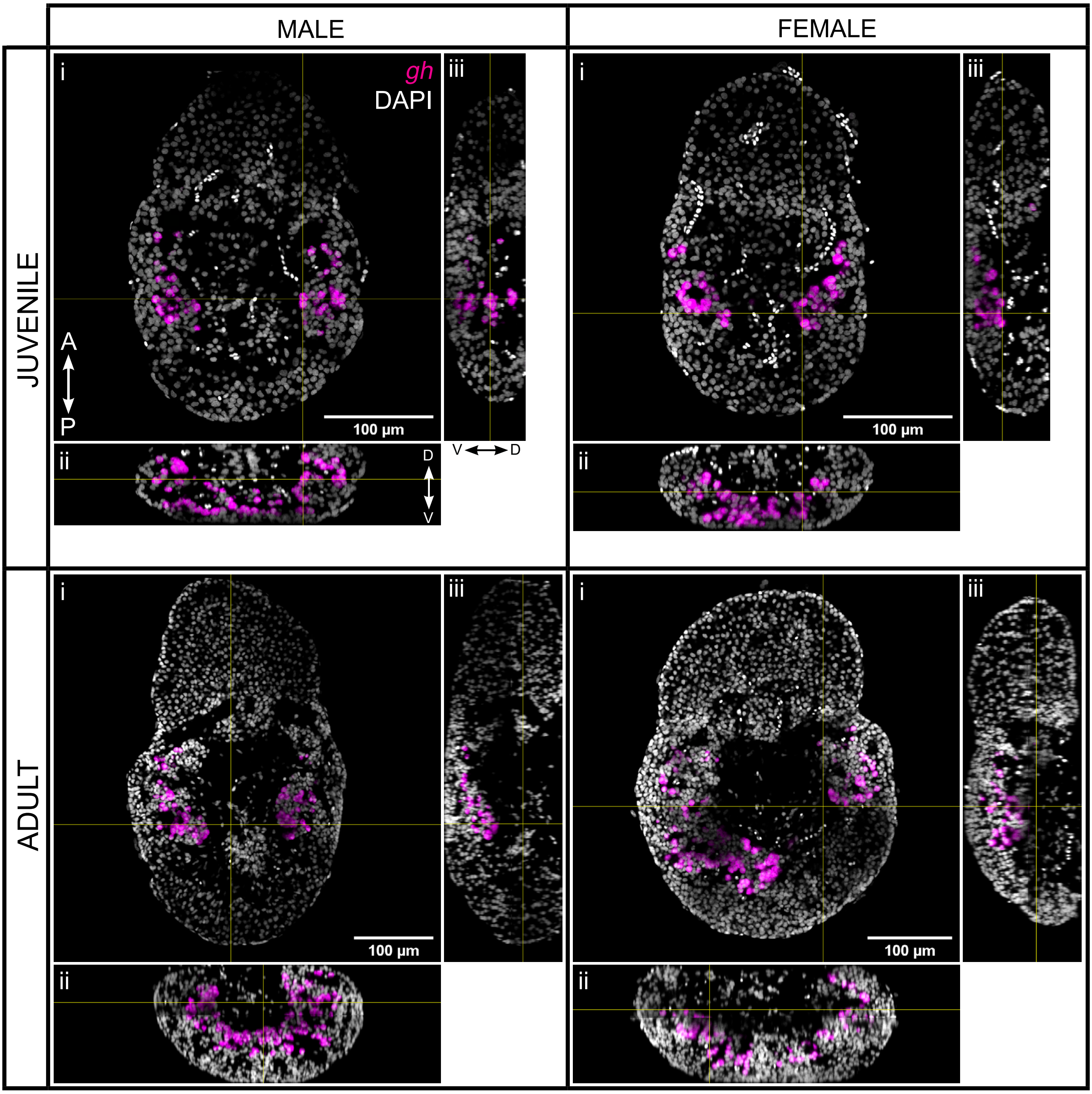


SUPP. FIG. 7

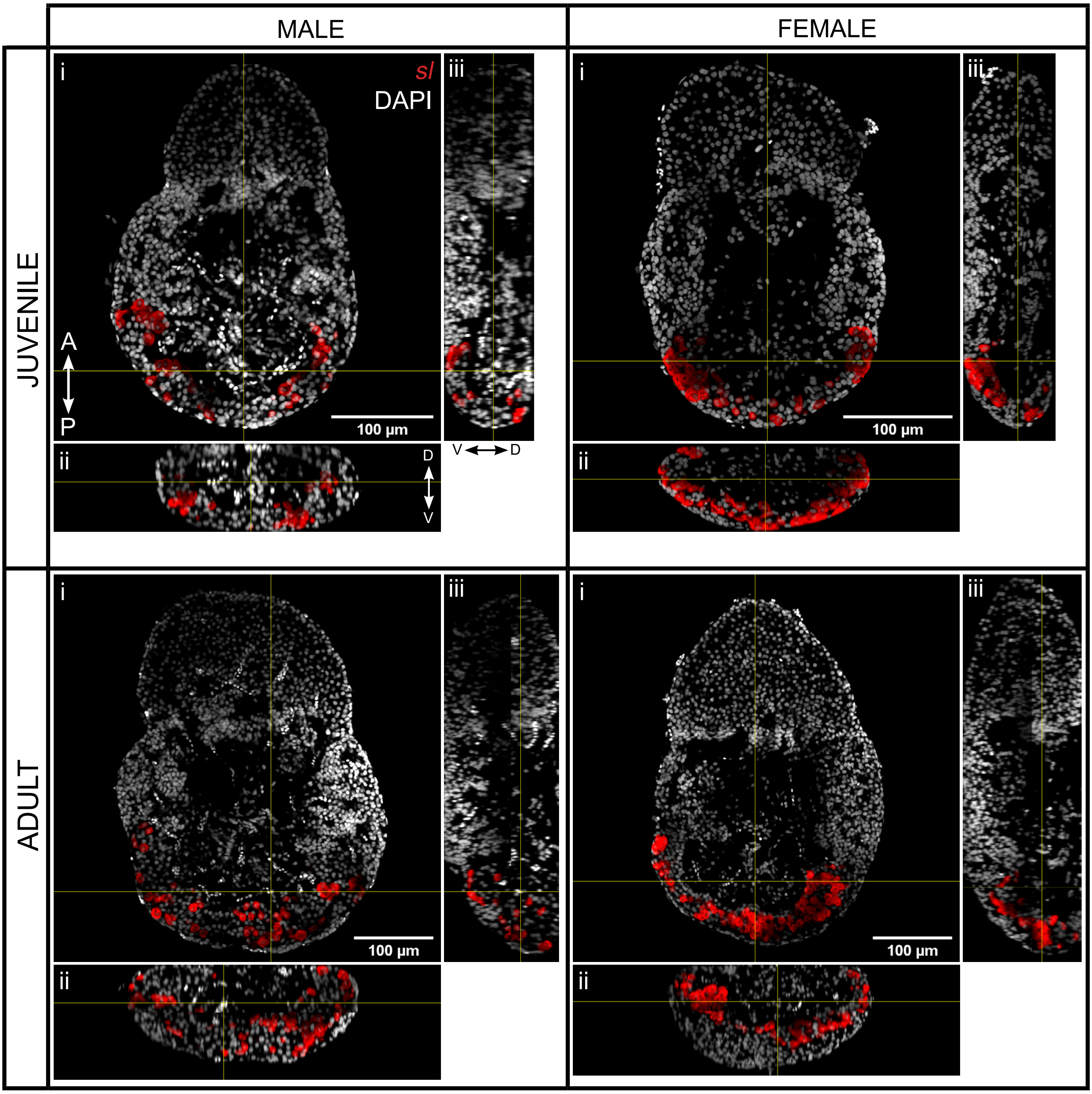


SUPP. FIG. 8

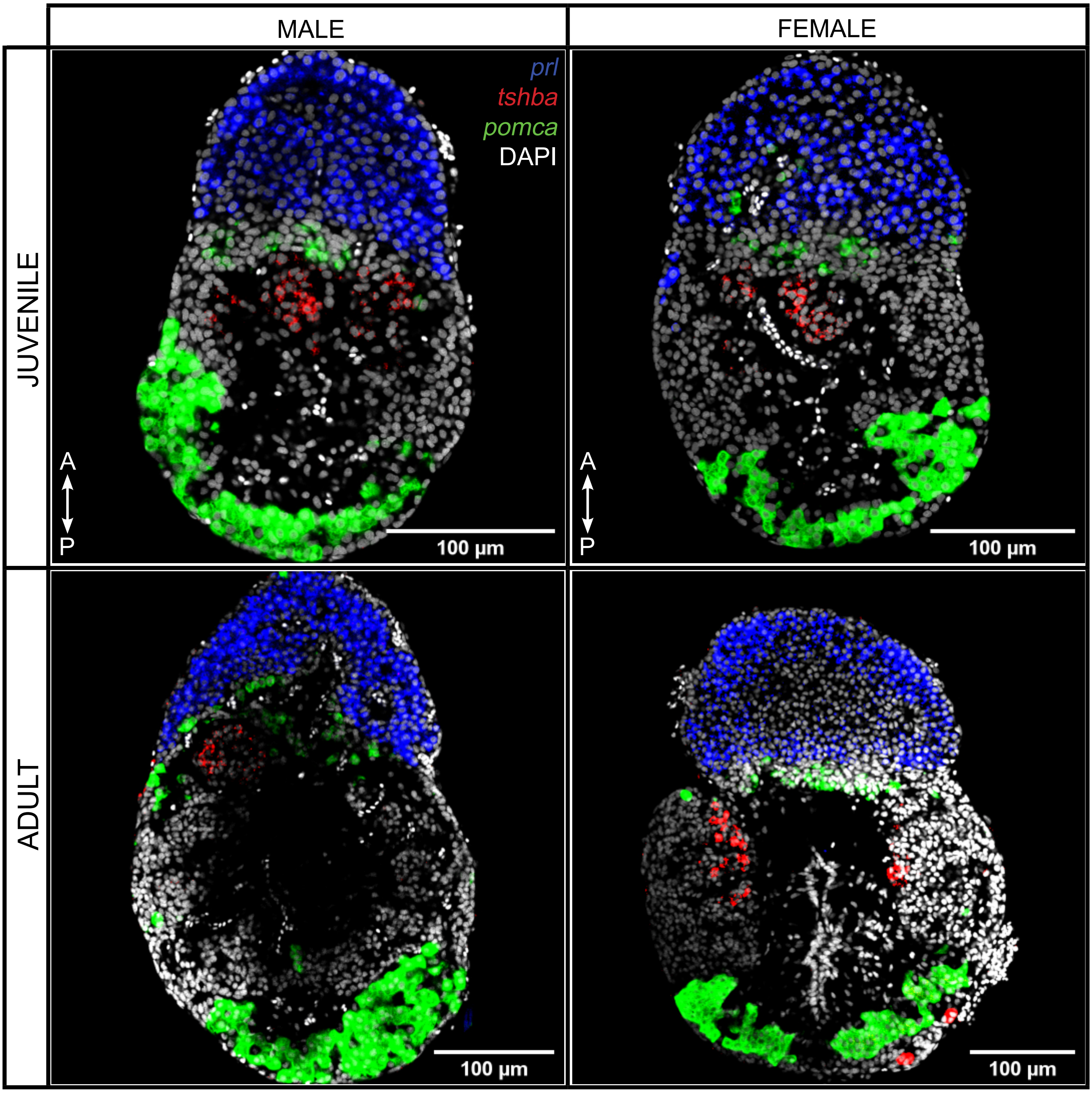
MALE
FEMALE
JUVENILE
ADULT
prl
tshba
pomca
DAPI
A
P
100 µm
A
P
100 µm
100 µm
100 µm

SUPP. FIG. 9

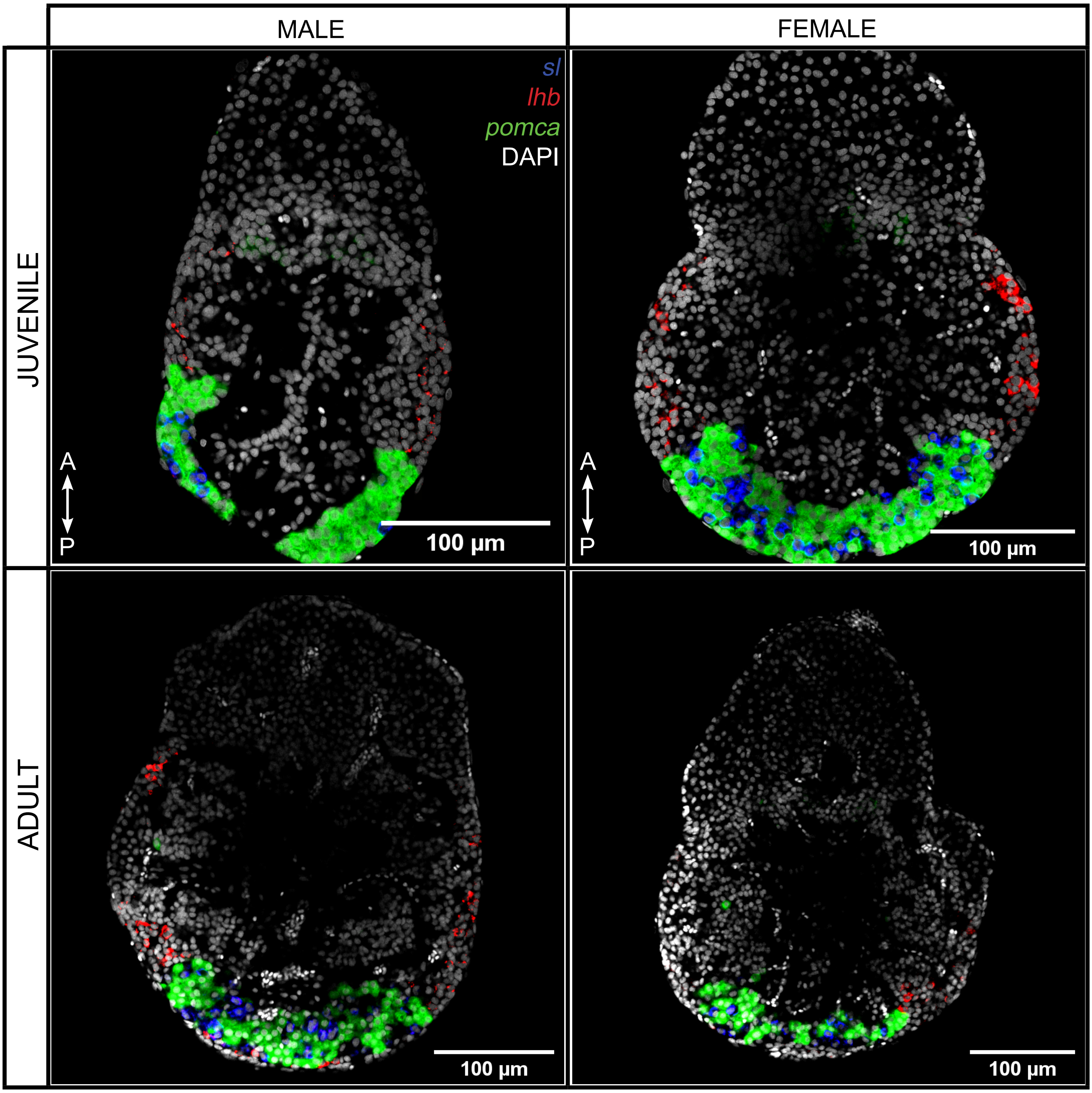
MALE
FEMALE
JUVENILE
ADULT
sl
lhb
pomca
DAPI
A
P
100 µm
A
P
100 µm
100 µm
100 µm

SUPP. FIG. 10

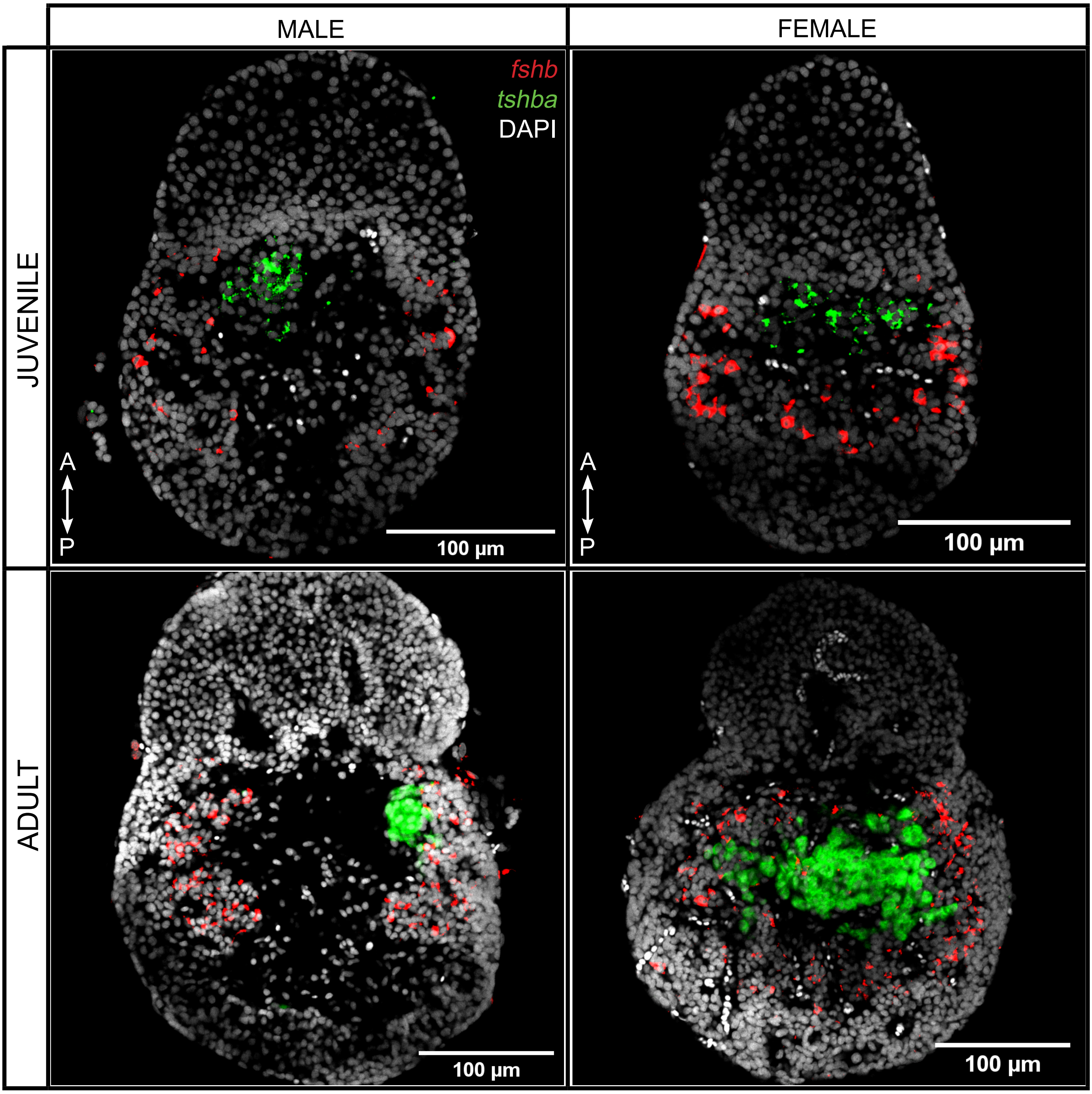
MALE
FEMALE
JUVENILE
ADULT
fshb
tshba
DAPI
A
P
A
P
100 µm
100 µm
100 µm
100 µm

SUPP. FIG. 11

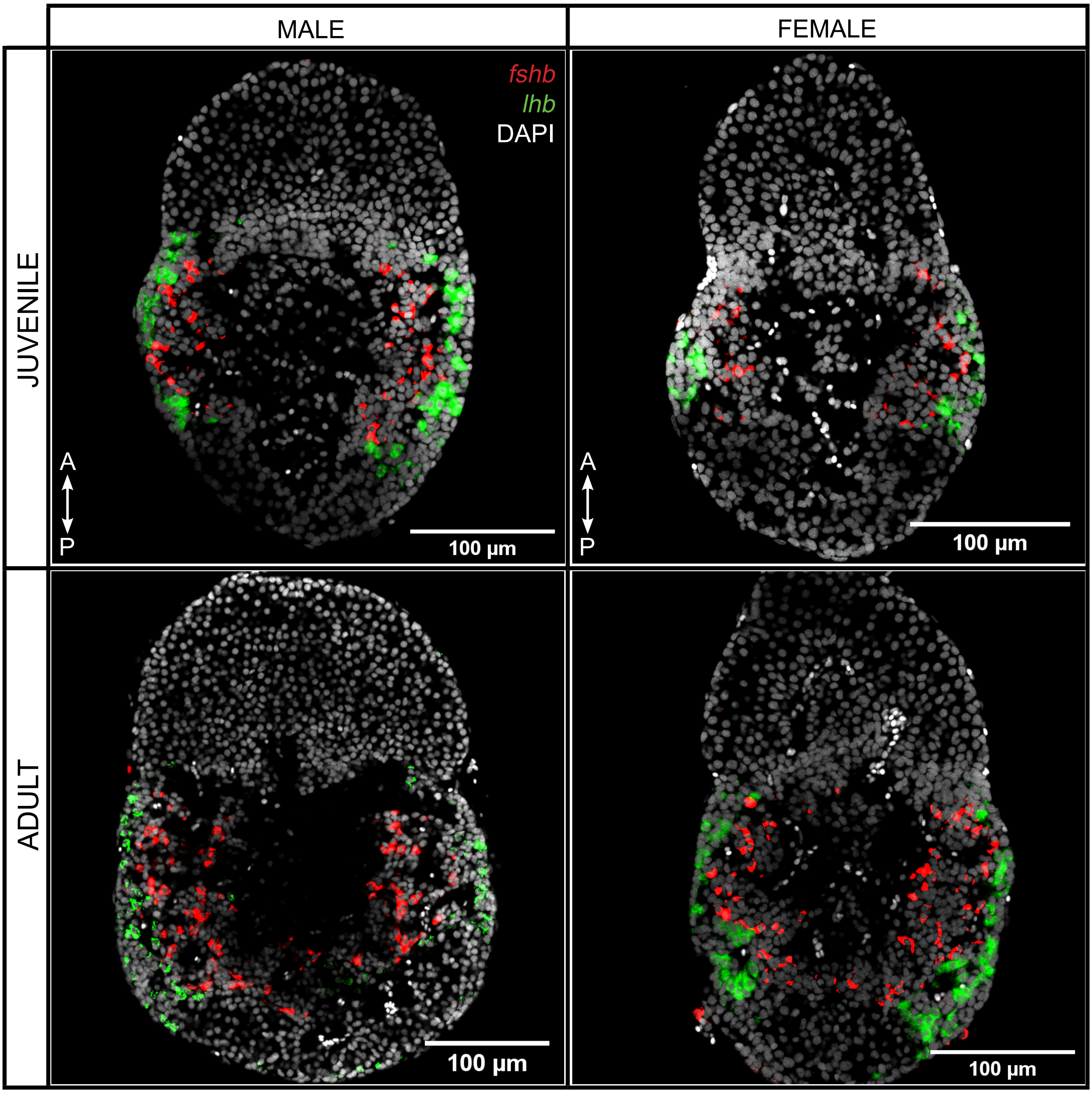
MALE
FEMALE
JUVENILE
ADULT
fshb
lhb
DAPI
A
P
A
P
100 µm
100 µm
100 µm
100 µm

SUPP. FIG. 12

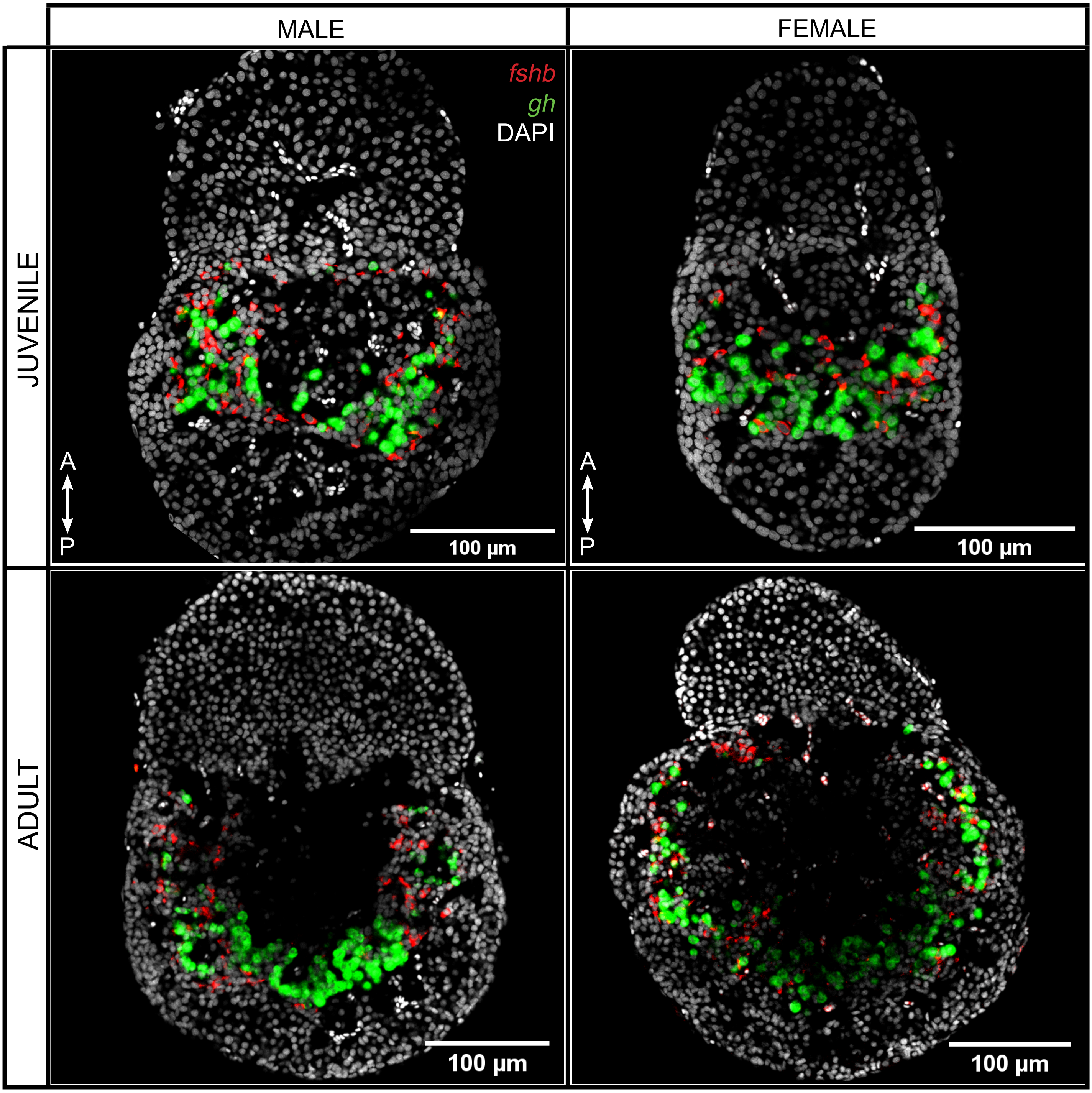

MALE
FEMALE
JUVENILE
ADULT
fshb
gh
DAPI
A
P
A
P
100 µm
100 µm
100 µm
100 µm


SUPP. FIG. 13

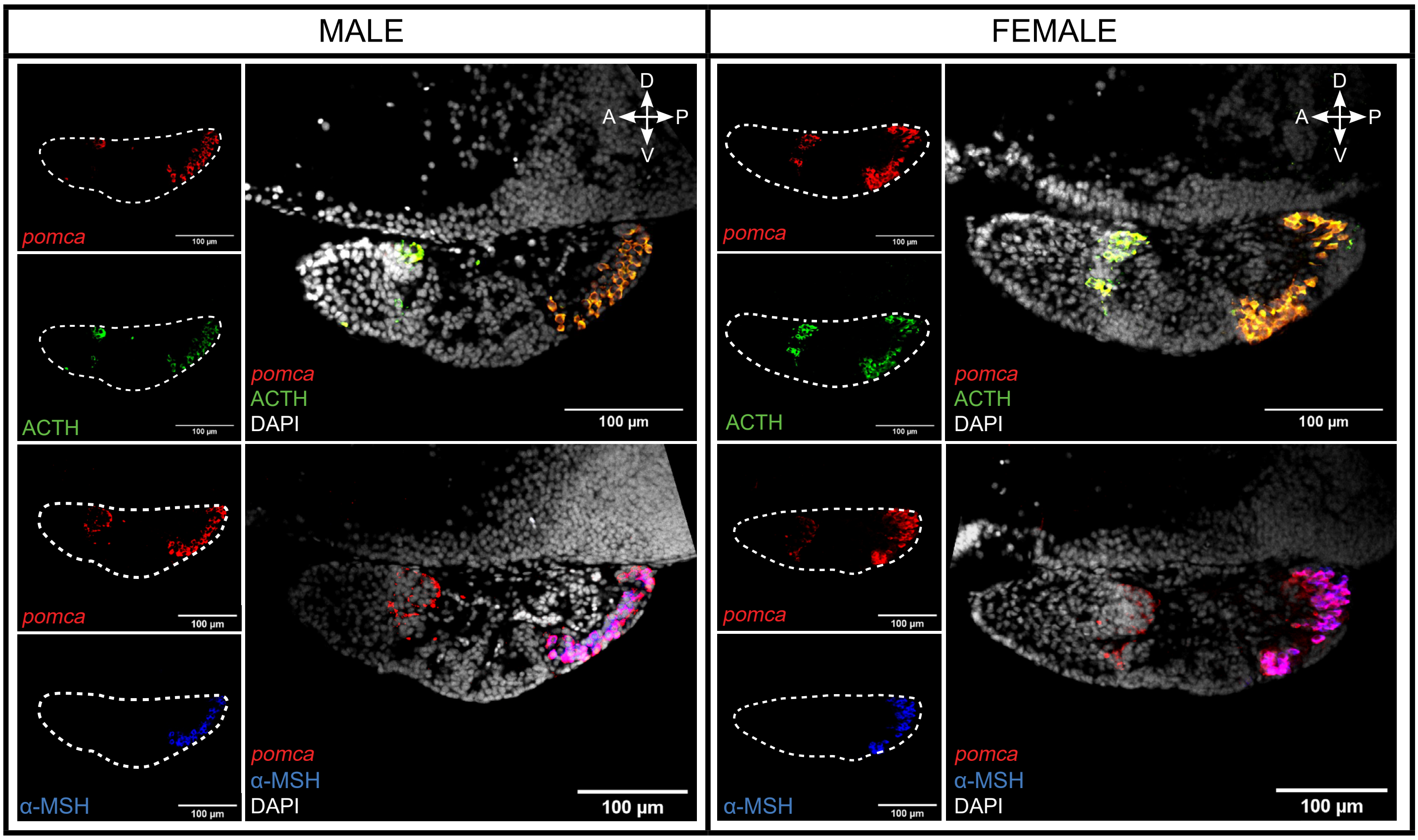


SUPP. FIG. 14
